# Reproducible Brain Charts: An open data resource for mapping brain development and its associations with mental health

**DOI:** 10.1101/2025.02.24.639850

**Authors:** G Shafiei, NB Esper, MS Hoffmann, L Ai, AA Chen, J Cluce, S Covitz, S Giavasis, C Lane, K Mehta, TM Moore, T Salo, TM Tapera, ME Calkins, S Colcombe, C Davatzikos, RE Gur, RC Gur, PM Pan, AP Jackowski, A Rokem, LA Rohde, RT Shinohara, N Tottenham, XN Zuo, M Cieslak, AR Franco, G Kiar, GA Salum, MP Milham, TD Satterthwaite

## Abstract

Major mental disorders are increasingly understood as disorders of brain development. Large and heterogeneous samples are required to define generalizable links between brain development and psychopathology. To this end, we introduce the Reproducible Brain Charts (RBC), an open data resource that integrates data from 5 large studies of brain development in youth from three continents (*N*=6,346; 45% Female). Confirmatory bifactor models were used to create harmonized psychiatric phenotypes that capture major dimensions of psychopathology. Following rigorous quality assurance, neuroimaging data were carefully curated and processed using consistent pipelines in a reproducible manner with DataLad, the Configurable Pipeline for the Analysis of Connectomes (C-PAC), and FreeSurfer. Initial analyses of RBC data emphasize the benefit of careful quality assurance and data harmonization in delineating developmental effects and associations with psychopathology. Critically, all RBC data – including harmonized psychiatric phenotypes, unprocessed images, and fully processed imaging derivatives – are openly shared without a data use agreement via the International Neuroimaging Data-sharing Initiative. Together, RBC facilitates large-scale, reproducible, and generalizable research in developmental and psychiatric neuroscience.

## Introduction

Mental disorders are increasingly understood as disorders of brain development (Kessler et al., 2012; Casey et al., 2014; Bethlehem et al., 2022). Neuroimaging studies of brain development have the potential to track healthy brain maturation and identify deviations linked to psychopathology. However, large and diverse samples are necessary to detect reliable patterns of neurodevelopment and identify generalizable links to psychopathology (Paus et al., 1999; Van Horn & Gazzaniga, 2002; Paus, 2010; Milham, 2012; Nooner et al., 2012; Marek et al., 2022; Zhou et al., 2023; Uddin et al., 2024; Brown et al., 2024). Multiple independent open science initiatives have facilitated this by sharing data publicly (Biswal et al., 2010; Milham et al., 2012; Nooner et al., 2012; Somerville et al., 2018; Howell et al., 2019; Satterthwaite et al., 2014; Alexander et al., 2017; Tobe et al., 2022; Volkow et al., 2018). However, while it is possible to aggregate data across independent studies (Biswal et al., 2010; Fair et al., 2013; Di Martino et al., 2013; Di Martino et al., 2017), it is not necessarily a straightforward process due to variation in neuroimaging and psychiatric phenotyping protocols used (Yan et al., 2013; Mennes et al., 2013). Obstacles in combining disparate data lead most investigators to use only a fraction of the data available (Milham, 2012; Poldrack & Poline, 2015; Gilmore et al., 2018). To address these challenges, we introduce the Reproducible Brain Charts (RBC) initiative: a large-scale, open data resource for the developing brain and psychiatry.

RBC addresses five major obstacles. First, there is abundant evidence that large, high-quality samples are essential for defining reliable associations between brain features and behavioral phenotypes (Van Horn & Gazzaniga, 2002; Paus, 2010; Milham, 2012; Nooner et al., 2012; Zuo et al., 2014; Milham et al., 2021; Marek et al., 2022; Xu et al., 2023). This is particularly challenging for studies of brain development, where large samples are also necessary to define a normative growth curve (Bethlehem et al., 2022). To yield generalizable results, samples must not only be large, but also diverse (Paus, 2010; Nooner et al., 2012; Volkow et al., 2018; Laird, 2021; Brown et al., 2024; Kang et al., 2024). Basic dimensions of diversity include age, sex, socioeconomic status, clinical diagnosis, race, and genetic ancestry (Paus, 2010; Nooner et al., 2012). Accordingly, large-scale samples must encompass such variability to identify generalizable patterns of neurodevelopment and their links to mental health. In response, RBC has assembled a diverse dataset from five major developmental cohort studies across three continents, spanning various recruitment strategies and thereby enriching both psychopathology and demographic diversity. This model serves as a foundational starting point for future expansions and the inclusion of data from additional studies and cohorts.

Second, combining psychiatric phenotypic data across large-scale studies of brain development presents multiple challenges (McElroy et al., 2020; Polanczyk et al., 2014; Luningham et al., 2020; Bauer and Hussong 2009; Curran and Hussong 2009). An initial challenge is obvious: different studies often employ disparate assessment tools to measure the same construct. Moreover, even when the same assessment is used, important psychometric properties may vary across populations. Aggregated data thus requires careful harmonization of both measures and response properties. RBC addresses these issues by mapping differing tools to a common framework using a bifactor model (Reise, 2012; Lahey et al., 2012; Caspi et al., 2014; Kotov et al., 2017, 2019; Hoffmann et al., 2022–2024). Bifactor models provide a robust solution by extracting a general factor (the “*p*-factor”) that captures shared variance across symptoms and distinct domain-specific factors (e.g., internalizing or externalizing symptoms) that remain independent of the general factor. This approach effectively summarizes correlated psychiatric symptoms and diagnoses, aligning with dimensional models like HiTOP (Kotov et al., 2017, 2019) and RDoC (Cuthbert & Insel, 2013).

A third major challenge is that both image acquisition parameters and image processing procedures vary considerably across large-scale studies. Even within multi-site studies with harmonized protocols, variation in scanners and protocol adherence introduces significant technical variability (Poldrack & Poline, 2015; Laird, 2021). Furthermore, many large-scale studies do not publicly release fully processed data. When they do, different studies utilize discrepant image processing pipelines, further complicating data aggregation and the reproducible integration of findings across studies (Carp, 2012; Baker, 2016; Nichols et al., 2017; Botvinik-Nezer et al., 2020; Bhagwat et al., 2021; Li et al., 2024). To address this, we processed all RBC data with standard tools like FreeSurfer (Fischl, 2012) and the Configurable Pipeline for Analysis of Connectomes (C-PAC; Craddock et al., 2013b). C-PAC’s highly configurable workflow allowed us to apply multiple versions of functional image processing uniformly across the entire data resource. Notably, all image processing steps were executed within the FAIRly-big framework (Wagner et al., 2022), which allowed us to maintain a detailed audit trail via DataLad (Halchenko et al., 2021). Fully documented and traceable data curation and processing enable researchers to rerun and adapt workflows for their analyses, facilitating harmonization and integration across datasets (Bellec et al., 2017). Following such reproducible image processing, we further reduced acquisition-related variation using statistical techniques adapted from computational genomics to harmonize imaging features (Johnson et al., 2007; Fortin et al., 2017; Fortin et al., 2018; Pomponio et al., 2020; Chen et al., 2022; Hu et al., 2023; Bridgeford et al., 2025).

Fourth, variation in data quality remains a very important confound in neuroimaging research, with in-scanner motion profoundly impacting imaging features like functional connectivity. This challenge is particularly acute for studies of brain development and psychiatry, as younger children and individuals with significant symptoms tend to move more during scanning sessions (Satterthwaite et al., 2012; Fair et al., 2013). Without meticulous quality control (QC), the substantial effects of data quality can easily overshadow the more subtle variations associated with brain development or psychopathology, leading to spurious associations that may be misinterpreted as biologically meaningful (Power et al., 2012; van Dijk et al., 2012; Satterthwaite et al., 2012; Yan et al., 2013; Murphy et al., 2013; Gilmore et al., 2021). To address this, RBC provides an extensive array of QC metrics alongside specific suggestions for exclusion criteria, bolstering the validity of the integrated data.

Fifth and finally, the mechanics of data access remain a major obstacle for investigators who hope to exploit large-scale data resources. Administratively cumbersome data use agreements (DUAs) often delay researchers and diminish the returns on public investments (Milham et al., 2018; White et al., 2020; Tedersoo et al., 2021; Laird, 2021; Jwa & Poldrack, 2022; Di Martino et al., 2013; Zuo et al., 2014). In contrast, the fully de-identified data in RBC is released as a completely open resource. This open-access approach allows all harmonized psychiatric phenotypes and imaging data to be freely shared via the International Neuroimaging Data-sharing Initiative (INDI; Mennes et al., 2013) without the need for a DUA, thereby accelerating research and maximizing the impact of public investments.

## Results

### RBC aggregates five diverse neurodevelopmental datasets

RBC integrates demographics, psychiatric phenotyping, and both T1-weighted structural and resting-state and task functional Magnetic Resonance Imaging (MRI) data from five diverse and prominent developmental cohorts (see **Table S1**; total *N* = 6,346). Specific studies include: (1) Brazilian High Risk Cohort (BHRC; Salum et al., 2014); (2) Developmental Chinese Color Nest Project (CCNP; Liu et al., 2021; Fan et al., 2023); (3) Healthy Brain Network (HBN; Alexander et al., 2017); (4) Nathan Kline Institute–Rockland Sample (NKI; Tobe et al., 2022); (5) Philadelphia Neurodevelopmental Cohort (PNC; Satterthwaite et al., 2014; Satterthwaite et al., 2016). The vast majority of data included in RBC is from childhood, adolescence, and early adulthood (i.e., age range: 5-22 years old), with the exception of the NKI dataset that also includes participants from across the lifespan (up to 85 years old). All datasets have a relatively balanced sex distribution (about 45% female across all datasets). In addition, RBC provides information about participant race and ethnicity, handedness, body mass index (BMI), participant education, parental education, and psychopathology (see below).

### RBC provides harmonized phenotypic data across studies

Selection of phenotypic instruments predated the RBC, and as such, reflects the design of each study. Specifically, the Child Behavior Checklist (CBCL) was used in BHRC, CCNP, HBN, and NKI while GOASSESS was used in PNC. The CBCL is a 120-item parent-report assessment of emotional and behavioral phenotypes (Achenbach & Rescorla, 2001). PNC did not include the CBCL, but rather assessed psychopathology using a highly-structured psychiatric screening interview (GOASSESS) (Calkins et al., 2015).

In RBC, we sought to derive major dimensions of psychopathology that both harmonized differences between samples assessed with the same instrument (i.e., the CBCL) as well as harmonizing differences between instruments (i.e., the CBCL vs. GOASSESS; see **Figure 1A**). To this end, we modeled psychopathology using a bifactor model (Reise, 2012; Lahey et al., 2012; Caspi et al., 2014). Specifically, bifactor models yield a general factor that represents overall psychopathology (also known as the “*p*-factor”) as well as orthogonal factors for more specific symptom domains (e.g., externalizing, internalizing, etc). We first evaluated 12 bifactor models based on previous literature to identify the model that best harmonized phenotypic data, minimizing between-study differences in CBCL (Hoffmann et al., 2022). Furthermore, we conducted an extensive evaluation of methods for harmonizing GOASSESS with CBCL using methods from item response theory and CFA (Hoffmann et al., 2023; Hoffmann et al., 2024). Ultimately, we developed a harmonized model that included a general psychopathology factor as well as specific factors for internalizing, externalizing, and attention symptoms (Hoffmann et al., 2023; **Figure 1B**). Model fits and factor structure for the item-level harmonized CBCL– GOASSESS model are available in **Tables S2–3**. Participant scores for the general psychopathology factor and each of the domain-specific factors are publicly shared in RBC.

**Figure 1.**
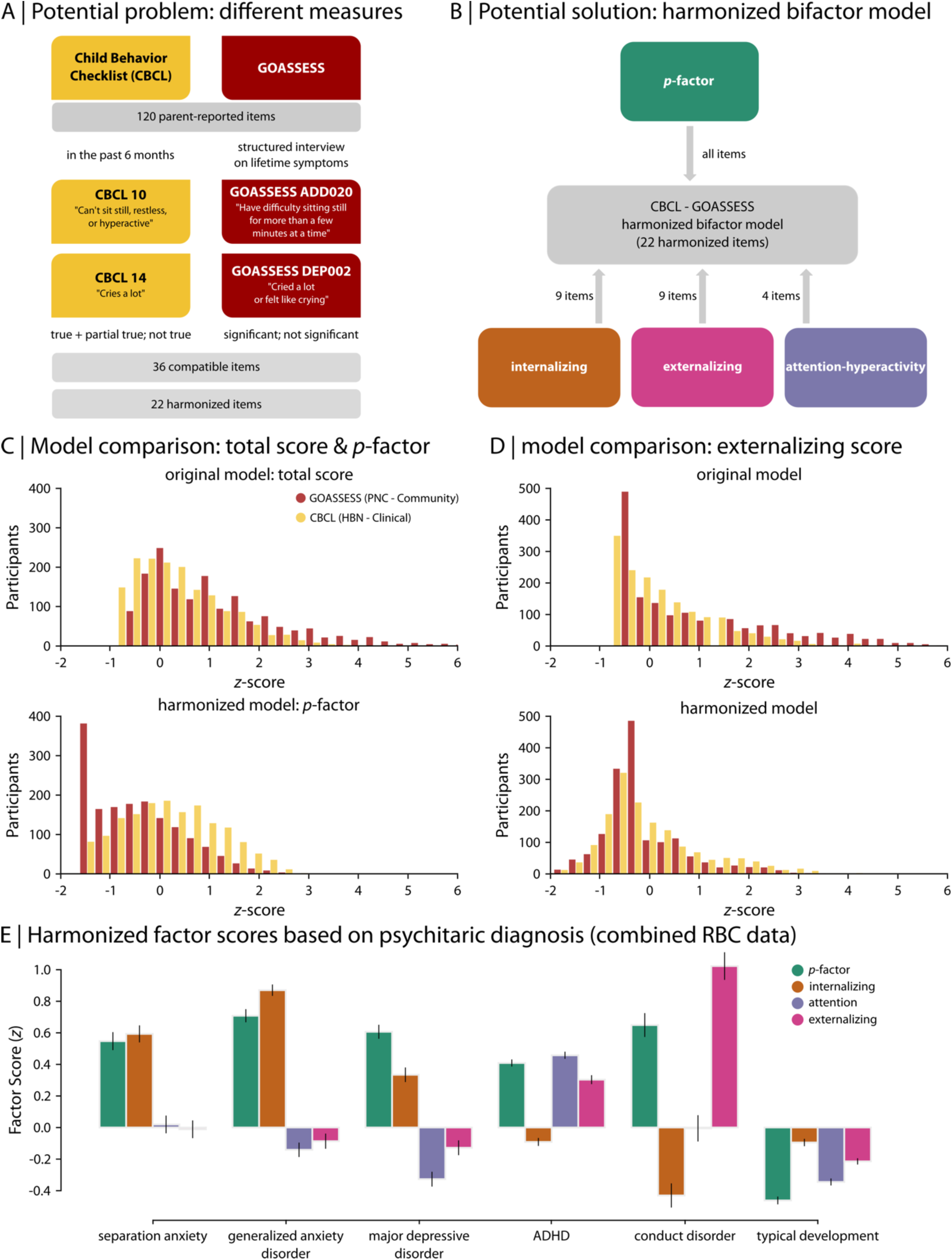
RBC provides harmonized phenotypic data. (A) Phenotypic data were separately collected by studies included in RBC using different instruments. Child Behavior Checklist (CBCL) was used in BHRC, CCNP, HBN, and NKI and GOASSESS was used in PNC. Both instruments contain a subset of variables that are conceptually overlapping, but it is not possible to directly compare them without phenotypic data harmonization. To harmonize phenotypic data, expert-based 1-to-1 semantic item-matching was used to identify 36 compatible items across studies, which were then reduced to 22 harmonized items. (B) The McElroy bifactor model was used to capture major dimensions of psychopathology based on the 22 harmonized items, yielding a factor that represents general psychopathology (i.e., *p*-factor) and orthogonal factors that represent specific domains (i.e., externalizing, internalizing, and attention factors). (C, D) The impact of the RBC phenotypic data harmonization is demonstrated for PNC (GOASSESS questionnaire) and HBN (CBCL checklist), depicting original total scores (i.e., normalized sum scores based on full item sets) and harmonized factors for *p*-factor (C) and externalizing factor (D). Given that HBN primarily includes help-seeking youth, higher levels of psychopathology are expected for HBN compared to PNC (a community-based sample). This difference is more evident in harmonized data. (E) The factor scores aligned with clinical diagnostic categories defined by DSM. For example, higher externalizing factor and *p*-factor values were observed in conduct disorder, while higher internalizing factor and *p*-factor values were observed in major depression and anxiety disorders. All general and specific factors were low in typical development.

An illustrative example of the impact of harmonization on psychiatric phenotypes in RBC is apparent when comparing the overall psychopathology factor (*p*-factor) and the externalizing factor in PNC and HBN (**Figure 1C-D**). As noted above, PNC and HBN used different instruments to assess psychiatric phenotypes (CBCL vs. GOASSESS). They also differed in their sample recruitment strategy: while PNC was a community-based sample, HBN primarily consists of help-seeking youth with significant symptoms of mental illness. As such, we expect higher levels of psychopathology in HBN than PNC. Notably, before harmonization, this difference was not apparent (**Figure 1C**; original psychopathology score: two-sample *t*-test: *t* = - 8.4e-15, *p* = 0.9). Following harmonization, higher levels of psychopathology were evident in HBN as expected (**Figure 1C**; harmonized psychopathology score: two-sample *t*-test: *t* = 22.9, *p* = 1.3e-110). These harmonized factor scores aligned with clinical diagnostic categories defined by the DSM (**Figure 1E**). For example, conduct disorder was marked by elevated externalizing factor and *p*-factor values, whereas major depression and generalized anxiety were marked by higher *p*-factor and internalizing factor scores. In contrast, all general and specific factors were low in typically developing youth. Together, this harmonization process allows for direct pooling of phenotypic data across the diverse studies included in RBC.

### Neuroimaging data are curated in a fully reproducible manner in RBC

We aggregated structural and functional neuroimaging data from multiple independently collected, large-scale data resources in RBC, while addressing challenges due to variations in imaging data acquisition (**Tables S4–5**). Imaging metadata, described by the Brain Imaging Data Structure (BIDS; Gorgolewski et al., 2017), can be used to automatically configure processing workflows (e.g., “BIDS Apps”; Gorgolewski et al., 2017). Such automatic configuration facilitates processing of large datasets that may have been collected using different data acquisition protocols. However, given that automatic configuration of processing workflows relies on metadata, any inaccuracies in metadata can lead to pipelines that are reproducible but wrong. This reliance makes careful curation of metadata essential as any issues at the curation step will influence all subsequent analysis. Yet, data curation is typically an ad-hoc process – often involving poorly-recorded manual intervention that is not reproducible, compromising the chain of reproducibility from the start.

To overcome these challenges, we converted all datasets to the BIDS format and curated the data using the reproducible workflows provided by the Curation of BIDS (CuBIDS; Covitz et al., 2022) package. CuBIDS summarizes the heterogeneity in image acquisition and facilitates metadata curation. Moreover, CuBIDS uses DataLad (Halchenko et al., 2021) to ensure reproducibility throughout the curation process. Each study in RBC was curated with CuBIDS, yielding summary tables that describe and summarize heterogeneity in image acquisition (**Data S1-5)**. For example, the PNC dataset had 3 different CuBIDS parameter groups for structural images (i.e., T1-weighted MRI scans), separating T1 images to a main group with the majority of scans (n=1597) and 2 variant groups with only a few scans each (e.g., n=3 and n=1). The CuBIDS summary files also indicate the source of the variance in T1 image acquisition parameters. For example, the sources of the variance in PNC T1 images were obliquity for one variant group and slightly different echo and repetition times for the other variant group.

Similarly, HBN had 10 different CuBIDS parameter groups for T1-weighted MRI scans. The 10 different parameter groups separated T1 images to 6 main groups and 4 variant groups based on data acquisition sites and protocols, consistent with the fact that HBN is a multi-site dataset and uses multiple acquisition protocols. From those 6 main groups, 4 dominant acquisition groups contained the majority of scans protocol at each site (i.e., n=1157 at RU site; n=911 at CBIC site; n=343 at SI site; n=88 at CUNY site) while 2 groups contained a smaller number of scans with a different acquisition protocol (i.e., n=57 at CBIC site; n=5 at CUNY site). Each of the 4 variant groups consisted of only a single scan based on slightly different acquisition parameters. Notably, functional MRI data were generally more heterogeneous, where the main source of the variance was the number of volumes acquired during fMRI scans. In all cases, our use of CuBIDS to curate the data in RBC bolstered our confidence in the metadata used by BIDS-apps to configure image processing pipelines.

### RBC provides fully-processed structural and functional MRI data

Differences in data processing pipelines across studies present significant barriers to aggregating data from multiple resources. To support cross-study analyses, we used uniform processing pipelines and maintained a comprehensive audit trail in RBC. Specifically, following data curation, we used consistent image processing pipelines across studies included to generate a standardized set of commonly used measures of brain structure and function (Van Essen et al., 2013; Glasser et al., 2016b; Bellec et al., 2017; Volkow et al., 2018; Somerville et al., 2018). To ensure reproducibility and transparency, we adopted the “FAIRly-big” framework (Wagner et al., 2022), which enabled all data preparation and analyses to be accompanied by a detailed audit trail in DataLad (Halchenko et al., 2021). This audit trail not only allows for tracking all steps in the processing pipeline but also provides a robust mechanism to rerun the pipelines, preserving methodological integrity. While the use of random numbers in imaging pipelines, such as FreeSurfer (Fischl, 2012), sMRIPrep (Esteban et al., 2024), and C-PAC (Craddock et al., 2013b), can lead to slight variations in derived outputs (e.g., zip files with differing shasums), the pipeline commands are fully rerunnable, and their execution history is carefully recorded. This ensures that the entire process remains transparent and auditable, even if certain imaging derivatives are not strictly identical upon re-execution. By combining consistent pipeline application with an exhaustive audit trail, the RBC framework provides both reproducibility and transparency that supports robust cross-study analyses of brain structure and function.

In RBC, we provide fully-processed data including an extensive set of commonly used structural and functional MRI data derivatives (**Figure 2**). Structural data derivatives include surface area, cortical thickness, gray matter volume, folding and curvature indices, as well as summary whole-brain measures such as total intracranial volume. Functional data derivatives include preprocessed time-series and functional connectivity matrices with multiple edge weights (quantified as pairwise Pearson and partial correlations between regional time-series; **Figure 2B**). In addition, RBC provides regional measures such as Regional Homogeneity (ReHo; Y. Zang et al., 2004), Amplitude of Low Frequency Fluctuation (ALFF; Zang et al., 2007), and fractional ALFF (fALFF; Zou et al., 2008) (**Figure 2C**). Both structural and functional data are available in parcellated format using 16 anatomical, functional, and multimodal atlases (see “Methods” for further details).

**Figure 2.**
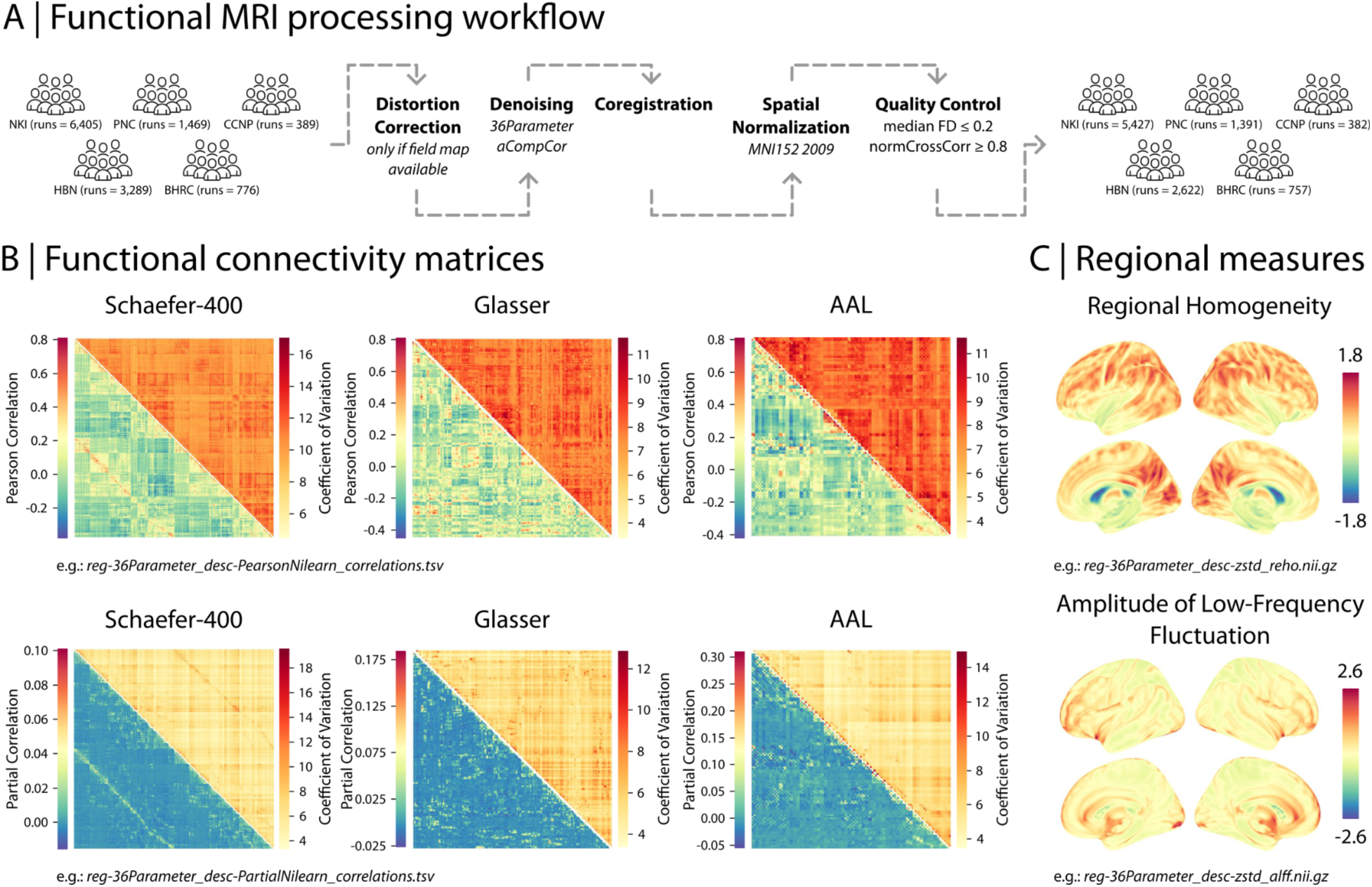
RBC provides processed neuroimaging data derivatives. (A) RBC integrates structural and functional neuroimaging data from five prominent neurodevelopmental datasets. Following careful data curation and consistent data preprocessing, neuroimaging data derivatives were estimated and publicly released in RBC. A few examples of group-average functional data derivatives are depicted: (B) functional connectivity matrices for 3 atlases with multiple edge weights (quantified as pairwise Pearson and partial correlations between processed regional time-series) and their corresponding coefficients of variation (calculated as the ratio between the standard deviation of the correlation values and the average absolute correlation value within each matrix); (C) regional functional properties such as Regional Homogeneity (ReHo) and Amplitude of Low Frequency Fluctuation (ALFF).

### RBC data is accompanied by harmonized measures of quality control (QC)

Data quality is one of the most important confounding factors in neuroimaging research. Image quality – driven mainly by in-scanner motion – affects both measures of brain structure (Reuter et al., 2015; Rosen et al., 2018; Gilmore et al., 2021) and functional connectivity (Power et al., 2012; van Dijk et al., 2012; Satterthwaite et al., 2012; Yan et al., 2013). The impact of data quality is even more pronounced in studies of development and psychopathology, where younger individuals and those with psychiatric symptoms tend to have higher in-scanner motion (Satterthwaite et al., 2012; Fair et al., 2013; Kong et al., 2014). Inconsistent QC criteria complicate cross-study comparability, yielding different samples and discrepant results from the same data. Thus, harmonized QC metrics that can be used for sample construction and model covariates (e.g., continuous QC metrics) are required to account for the impact of data quality.

To ensure consistent sample selection and comparable cross-study analyses, we generated harmonized measures of neuroimaging data QC in RBC. These measures are accompanied by specific QC guidelines that allow for consistent quality assurance. Summary information on the number of participants with structural and functional scans before and after applying RBC’s recommended QC is available in **Figures S1–S5**. Overall, approximately 90% of RBC data had adequate structural and functional QC (range: 75% for HBN to 99% for CCNP). Study- and modality-specific data on how RBC’s recommended QC affects sample selection procedures and data quantity are detailed in **Table S6**. As expected, image quality varied by study, with lower quality in studies with younger participants and greater psychopathology (e.g., HBN).

#### Structural MRI

RBC provides both harmonized QC information for structural MRI (sMRI) based on expert manual ratings as well as automated, quantitative indices of image quality. Specifically, every structural image was manually evaluated by 2-5 expert raters in multiple phases using Swipes for Science, a web application for binary image classification (Keshavan et al., 2019) (**Figure 3A**). Expert raters assigned ratings of “Pass” or “Fail” to four slices per each image. Each image was assigned a score per rater by averaging across slices, and then an overall QC score of “Pass”, “Artifact”, or “Fail” was assigned to the image based on the average rating across raters. Specifically, “Pass” or “Fail” labels were assigned to a given image if all raters were in agreement about its quality (i.e., all raters assigned the same label of either “Pass” or “Fail” to the image). The intermediate “Artifact” label was assigned to an image when there was disagreement between raters. The reliability of ratings was assessed across raters using a subset of images. We performed a rater reliability study for all six raters with 1,896 participants, totaling 7,584 slice ratings. The study showed a high level of agreement among raters (71.5% agreement; average Cohen’s Kappa = 0.71; **Figure 3B**). The majority of structural scans across studies included in RBC were labeled as “Pass” and were considered of adequate quality (82.6% on average across studies; **Figure 3C**). However, there was heterogeneity in data quality across datasets: HBN had the most “Artifact” and “Fail” scans compared to the other four studies, consistent with the fact that HBN includes younger help-seeking individuals with higher levels of psychopathology (with ADHD being the most frequent diagnosis). Furthermore, we found a positive trend between the average manual ratings and participant age (for age range of 5-21 years old; Spearman rho = 0.28; **Figure 3E**), consistent with lower in-scanner motion and improved image quality in older adolescents and younger adults.

**Figure 3.**
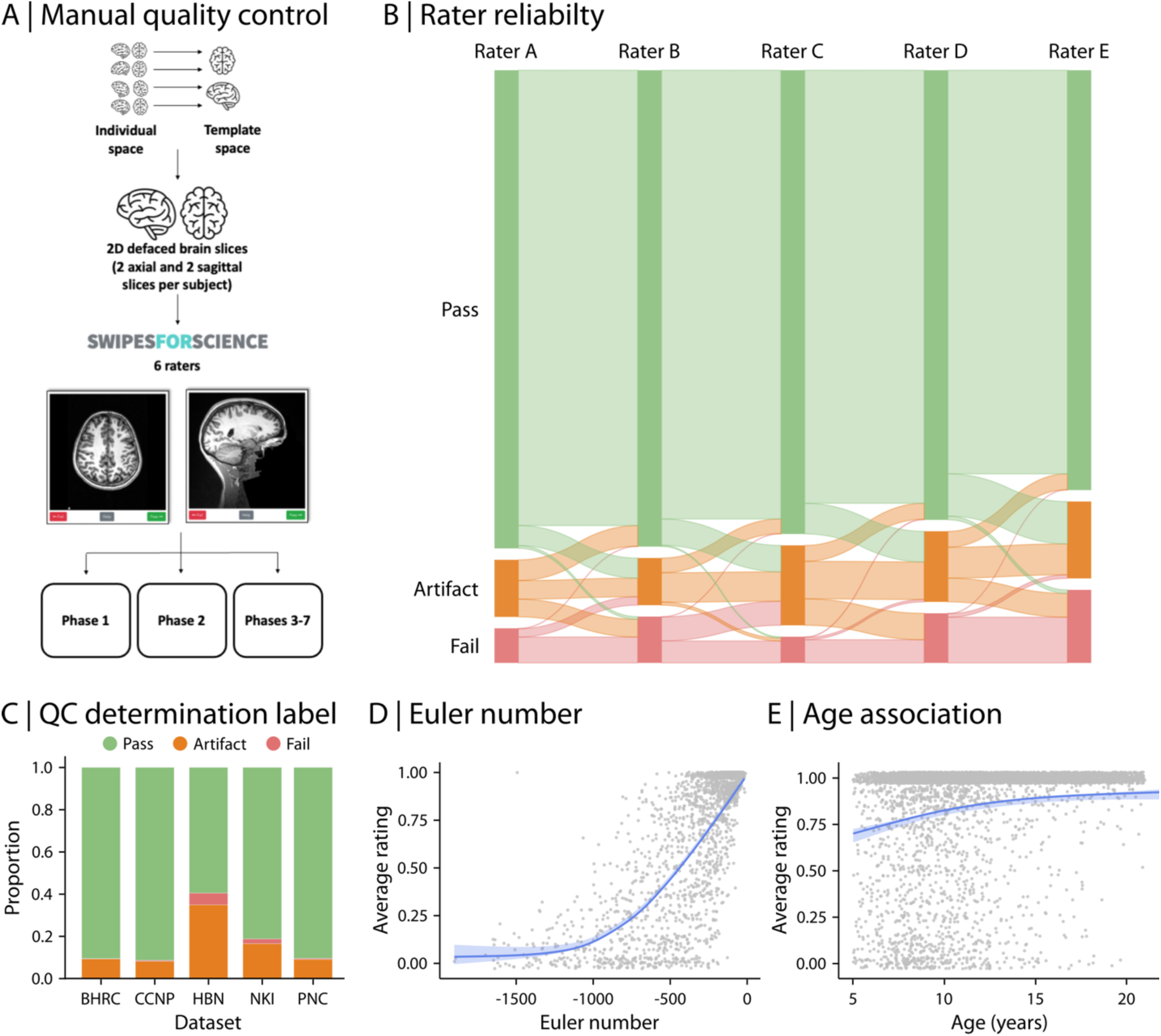
RBC data is accompanied by harmonized structural QC metrics. Harmonized structural QC metrics were estimated based on expert manual ratings and automated methods. (A) Each structural image was manually evaluated by 2-5 expert raters in 7 phases using Swipes for Science. 4 two-dimensional slices per participant (two axial and two sagittal slices extracted per structural scan) were created to rate each scan. Expert raters assigned ratings of “Pass” or “Fail” to four slices per each image. Each image was assigned a score per rater by averaging across slices, and then an overall QC score of “Pass”, “Artifact”, or “Fail” was assigned to the image based on the average rating across raters. Specifically, “Pass” or “Fail” labels were assigned to a given image if all raters were in agreement about its quality. The “Artifact” label was assigned to an image when there was any disagreement between raters. (B) Comparing the final QC categorization labels to expert ratings demonstrated that there was a high level of agreement among raters (71.5% agreement; average pairwise Cohen’s Kappa = 0.71). (C) The proportion of data with “Pass”, “Artifact”, and “Fail” labels are depicted for each study. HBN had the most “Artifact” and “Fail” scans compared to the other four studies, consistent with the fact that HBN includes help-seeking individuals with higher levels of psychopathology. (D) A continuous automated measure of data quality, Euler number, was estimated as part of the FreeSurfer processing pipeline. Euler number reflects the topological complexity of the reconstructed cortical surface, capturing variation in data quality within the coarse categories provided by manual ratings. As expected, Euler number was positively associated with the average manual expert ratings (Spearman rho = 0.63). (E) The average expert manual ratings was also positively associated with participant age, indicating higher in-scanner motion, and therefore, lower image quality in children and younger adolescents compared to older adolescents and younger adults (Spearman rho = 0.28).

In addition to the manual expert ratings of each T1w image, we calculated the Euler number for each structural scan. The Euler number is calculated as part of the FreeSurfer processing pipeline and reflects the topological complexity of the reconstructed cortical surface; the Euler number is lower when there are holes in the reconstructed cortical surface (i.e., more defects in surface reconstruction), and thus a ceiling of 0 indicates the highest possible quality by this metric. Previous reports have demonstrated that the Euler number is an accurate, fully-automated measure of data quality, capturing variation in data quality within the coarse categories provided by manual ratings (Rosen et al., 2018; Klapwijk et al., 2019; Elyounssi et al., 2022; Elyounssi et al., 2023). As expected based on prior work, we found that the Euler number was positively associated with the average manual expert ratings in RBC (Spearman rho = 0.63; **Figure 3D**). Thus, we recommend including Euler number as a covariate in secondary analysis of RBC data in addition to excluding data that do not meet RBC’s data quality criteria based on the categorical QC labels.

#### Functional MRI

fMRI derivatives in RBC are accompanied by extensive measures of QC, including multiple indices of both in-scanner motion and image registration quality (e.g., to the T1-weighted image or MNI template). To facilitate consistent QC across datasets, we generated a summary functional QC score. Specifically, fMRI runs with low in-scanner motion (i.e., median Framewise Displacement (FD) <= 0.2) and high image normalization quality (i.e., normalized cross correlation >= 0.8) were considered of adequate quality.

Using this approach, a relatively small proportion of participants per study were excluded due to poor data quality (9.7% excluded across studies). As expected, due to younger participants and higher prevalence of psychopathology, HBN had the largest number of individuals with increased in-scanner motion (**Figure 4A** & **B**). Furthermore, in-scanner motion decreased with age (Spearman rho = −0.22; **Figure 4C**). Median FD was positively associated with higher general psychopathology (*p*-factor; Spearman rho = 0.15; **Figure 4D**), suggesting that individuals with higher overall psychopathology tend to have higher in-scanner motion. Finally, median FD during fMRI was negatively related to manual ratings of structural image quality (Spearman rho = −0.45; **Figure 4E**), indicating that individuals with higher in-scanner motion during functional scans also had lower structural scan quality. Taken together, the extensive structural and functional QC metrics we provide allow for consistent quality assurance of imaging data in RBC. We strongly encourage researchers to use our recommended quality assurance ratings when working with RBC dataset. However, we also provide all data – even the scans that do not pass QC – to help accelerate progress on image QC research (see “Methods” for details on different versions of publicly-released RBC data).

**Figure 4.**
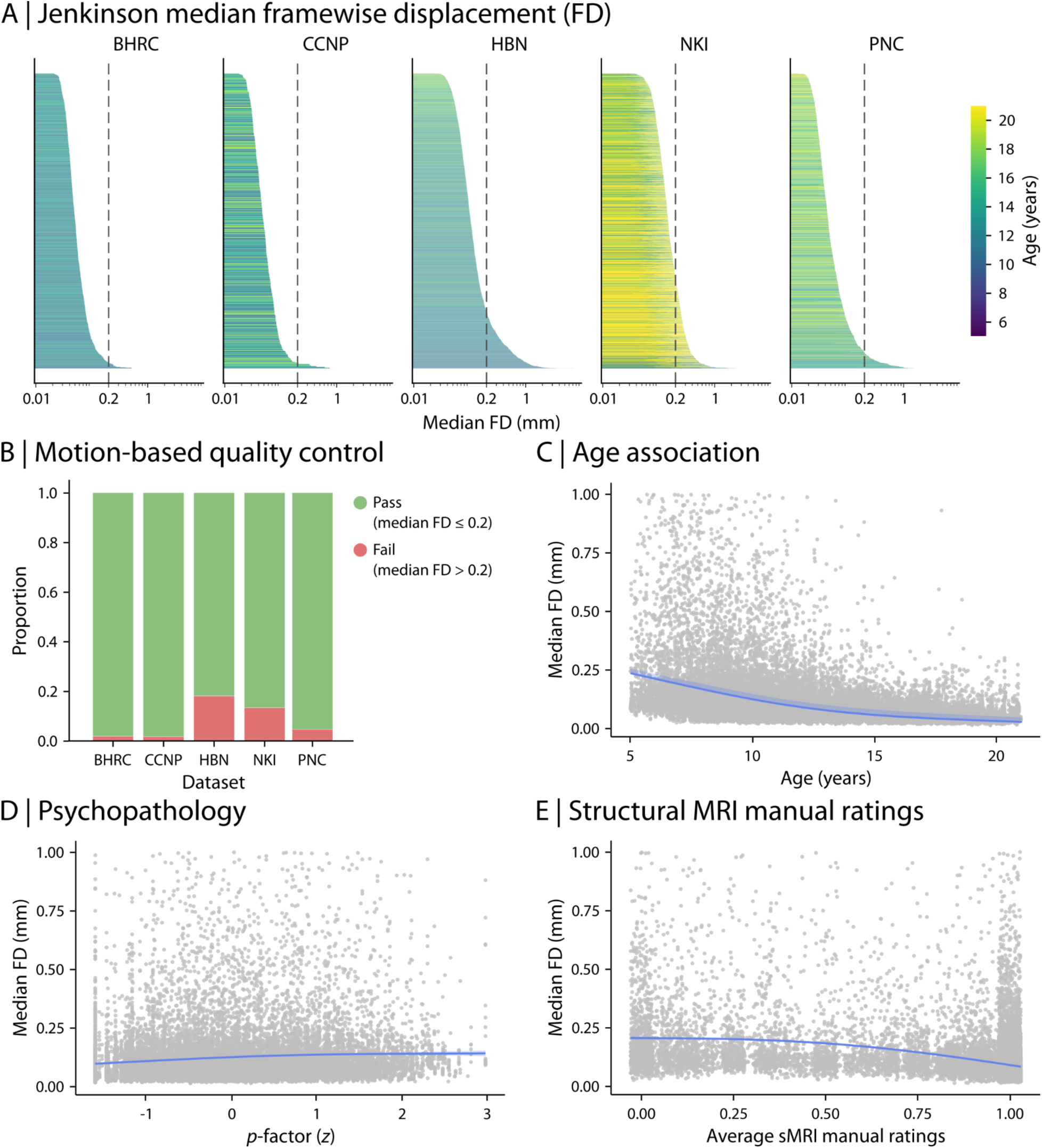
RBC data is accompanied by harmonized functional QC metrics. An extensive list of harmonized QC metrics were generated for functional RBC data, including summary measures of in-scanner motion, functional image to structural image registration (e.g., to T1- weighted scan or MNI template) and normalization quality indices. (A) A measure of in-scanner motion quantified as median Framewise Displacement (FD) is depicted as an example functional QC metric for all studies included in RBC. The dashed line indicates RBC’s recommended threshold for median FD, demonstrating that only a small proportion of individuals had high in-scanner motion, and hence, should be excluded from subsequent analysis of RBC data. The bar plots are color coded based on participants’ age. (B) As demonstrated in panel (A), the majority of RBC functional data pass the in-scanner motion threshold (i.e., median FD) of 0.2 across all studies. Consistent with the findings for structural QC (**Figure 3**), HBN had the largest number of individuals with high in-scanner motion. (C) Median FD was negatively associated with participant age, suggesting that older adolescents and younger adults tend to demonstrate lower in-scanner motion compared to children and younger adolescents (Spearman rho = −0.22). (D) In-scanner motion displayed a positive trend with general psychopathology (i.e., *p*-factor), suggesting that individuals with higher overall psychopathology tend to have higher in-scanner motion (Spearman rho = 0.15). (E) Median FD estimated from functional data was negatively associated with average manual ratings from structural data, indicating that individuals with higher in-scanner motion during functional scans also had lower structural scan quality (Spearman rho = −0.45).

### Neuroimaging features are associated with age and psychopathology in youth

To illustrate the utility of RBC, we examined how neuroimaging features were related to participant age and overall psychopathology. To ensure sensitivity to both linear and non-linear relationships, we used Generalized Additive Models (GAMs), while controlling for covariates such as sex and data quality (Pomponio et al., 2020; Sydnor et al., 2023; Luo et al., 2024). Prior to statistical analyses, imaging measures were harmonized using CovBat-GAM while protecting the effects of model covariates such as age (as a smooth term), sex, data quality, and *p*-factor (as linear terms) (Johnson et al., 2007; Fortin et al., 2017; Fortin et al., 2018; Pomponio et al., 2020; Chen et al., 2022; see **Figure S6** for site variation before and after harmonization). Sample sizes varied (*N* = 3,847 to 4,827) depending on data modality (i.e., structural versus functional features), analysis type (i.e., age versus psychopathology analysis), and whether QC was implemented; see **Table S7** for details.

#### Structural neuroimaging features are associated with development and psychopathology

We first assessed developmental associations with brain structure using structural MRI derivatives, including cortical thickness (CT), surface area (SA), and gray matter volume (GMV). The data were parcellated into 400 regions using the Schaefer-400 atlas (Schaefer et al., 2018). As a baseline, we evaluated age effects in combined RBC data without QC implementation or imaging data harmonization (**Figure 5A**). We found that developmental effects varied markedly between studies. For example, mean CT displayed a sharp decline with age in BHRC and PNC whereas these effects were more gradual in CCNP, HBN, and NKI. However, following QC and harmonization, the developmental patterns converged across studies (**Figure 5B**). Mean CT declined markedly with age; GMV and SA showed more gradual changes (CT and SA in **Figure 5** and GMV in **Figure S7**). Regional analyses identified heterogenous developmental effects in CT and SA across the cortex. CT decreased with age in most cortical regions, with the most prominent decrease observed in medial parietal, lateral and medial prefrontal, and temporal regions. SA displayed a combination of increases and decreases with age, such that medial frontal and parts of visual and motor cortices displayed an increase in SA whereas lateral parietal, temporal and prefrontal cortices displayed a decrease in SA during development (**Figure 5B**). Regional variations were consistent with the original analyses after controlling for global effects by including mean CT, SA, or GMV as model covariates in addition to other covariates (i.e., sex and Euler number; **Figure S8**). In addition, we repeated the analyses after implementing QC but before harmonizing imaging data (**Figure S9A**). We found that although QC removed data with low quality, data harmonization was required to effectively account for site and study differences.

**Figure 5.**
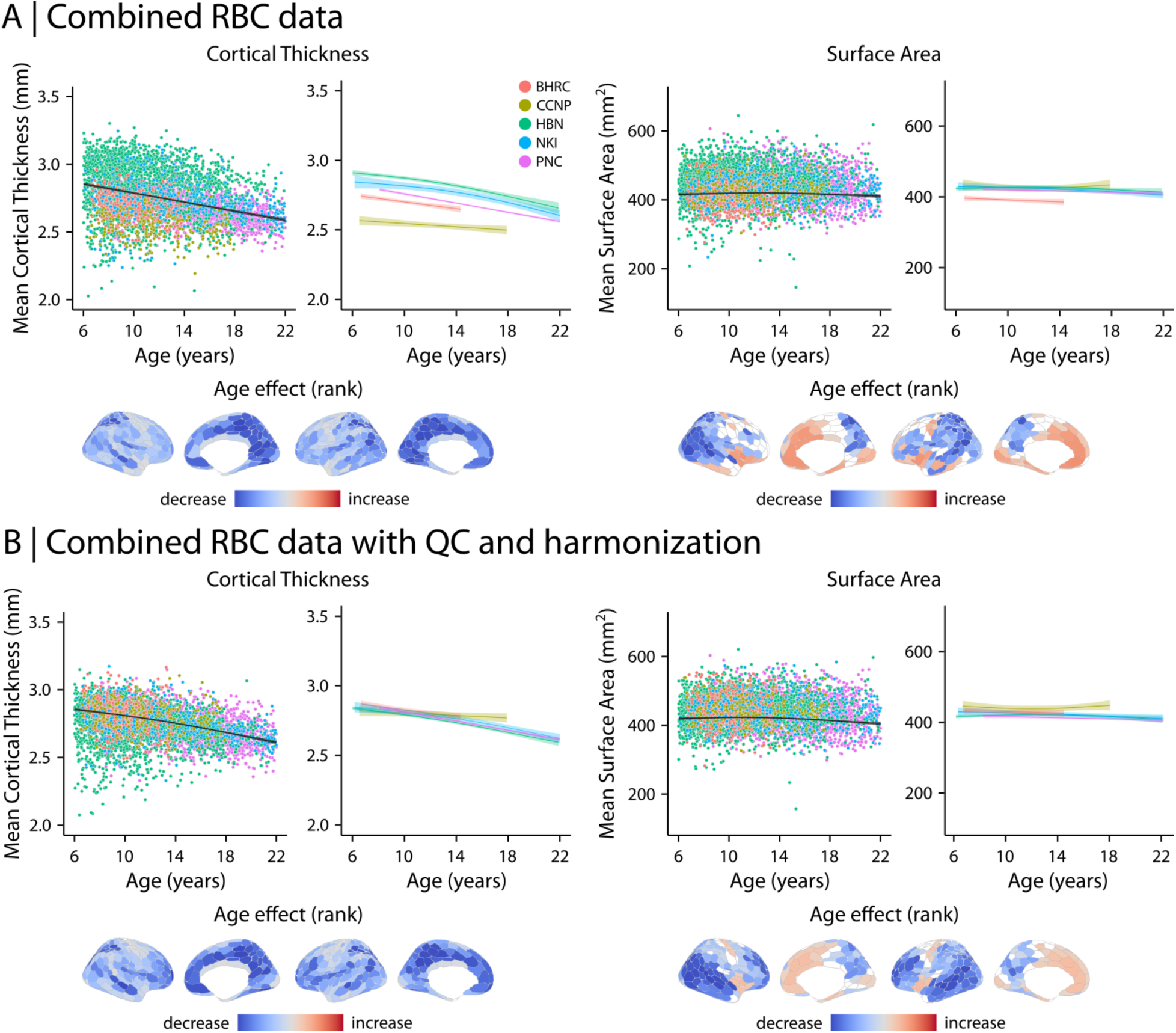
Structural data derivatives are associated with age in youth. Generalized Additive Models (GAMs) were used to examine the relationship between participant age and structural data derivatives, including cortical thickness (CT) and surface area (SA). (A) We used aggregated RBC data without QC implementation or neuroimaging data harmonization. We found an overall decrease in mean CT during development (with increasing age) while mean SA remained mostly unchanged. However, developmental effects varied markedly between studies as demonstrated by study-specific model fits (depicted with color coded curves). We also examined regional age effects using separate models for each brain region. Age effects were quantified using ranked partial *R*^2^ and are depicted on the cortical surface after correcting for multiple comparisons (FDR-corrected). Regional analysis demonstrated that CT decreases in most brain regions while SA decreases in lateral parietal cortices and increases in medial frontal areas. (B) To assess the effects of QC and data harmonization, we repeated the analysis after excluding data with “Fail” structural QC determination score and harmonizing neuroimaging data using CovBat-GAM. Following QC and data harmonization, the developmental effects on mean CT and mean SA became more similar and converged between studies as evident by study-specific model fits. Similar to before, the results demonstrated an overall decrease in mean CT with development. Regional analyses identified heterogeneous developmental effects on the cortex. CT decreased in the majority of brain regions (more prominent decrease in medial parietal, lateral and medial prefrontal, and temporal regions) while SA decreased in lateral parietal, temporal and prefrontal cortices and increased in medial frontal and visual and motor cortices.

We next examined the relationship between brain structure and psychopathology (*p*-factor) using the aggregated data (**Figure 6**). Similar to age-related findings, whole-brain associations between structural features and *p*-factor converged following QC and data harmonization (**Figure 6B**). Although regional analyses identified significant association with *p*-factor in both CT and SA in combined RBC data without QC and harmonization (**Figure 6A**), only SA was significantly associated with *p*-factor following QC and harmonization (**Figure 6B**). The results demonstrated that SA decreased with overall psychopathology, with the most prominent decrease in medial and lateral prefrontal cortices. We also repeated the analyses after implementing QC but before harmonizing imaging data (**Figure S9B**). This analysis suggested the presence of significant associations between CT and *p*-factor without data harmonization. However, these effects largely disappeared after harmonization with CovBat-GAM. Together, these findings underscore the importance of QC and harmonization when combining heterogeneous data. Importantly, including low quality data in the analysis or combining multi-site datasets without data harmonization may result in spurious associations between psychopathology and brain structure.

**Figure 6.**
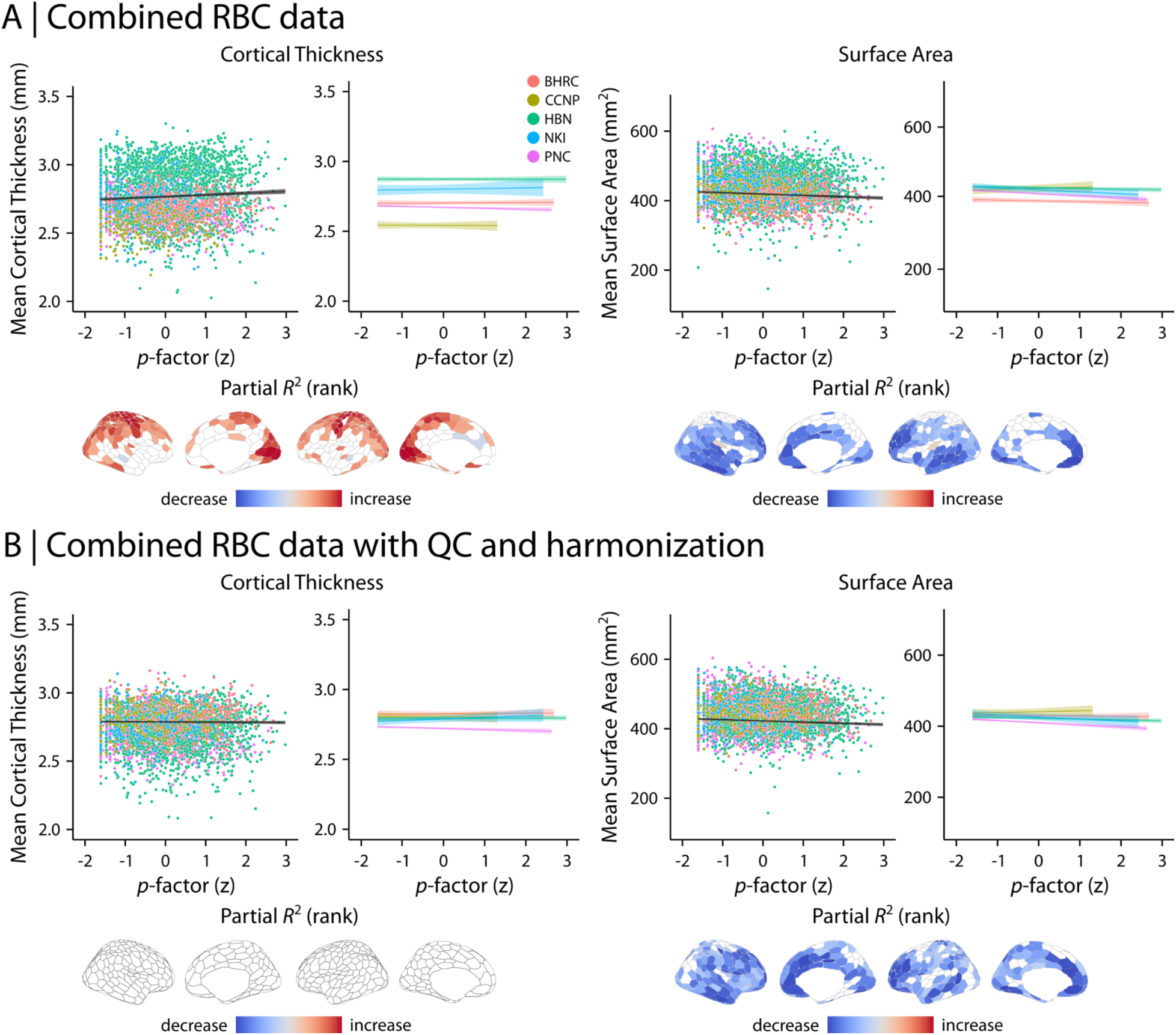
Structural data derivatives are associated with psychopathology in youth. Generalized Additive Models (GAMs) were used to examine the relationship between general psychopathology (i.e., *p*-factor) and structural data derivatives, including cortical thickness (CT) and surface area (SA). (A) GAM analyses using aggregated RBC data without QC implementation or neuroimaging data harmonization identified an overall increase in mean CT with increased *p*-factor and decreasing trend in mean SA with increased *p*-factor. Similar to age-effects (**Figure 5**), whole-brain associations between brain structure and psychopathology varied between studies. Regional analyses identified significant associations between structural features and *p*-factor in a subset of brain regions. (B) Following QC and neuroimaging data harmonization with CovBat-GAM, whole-brain association converged between studies, demonstrating no relationship between mean CT and *p*-factor and a decreasing trend in mean SA with increased *p*-factor. Regional analyses identified no significant associations between regional CT and psychopathology and significant decreasing patterns in SA with increased psychopathology (most prominent in medial and lateral prefrontal cortices). Note that CT was significantly associated with *p*-factor in a subset of regions in aggregated data before QC and data harmonization. This suggests including low quality data or combining multi-site datasets without data harmonization may result in spurious associations between brain structure and psychopathology.

#### Functional neuroimaging markers of age and psychopathology

To assess the developmental variations in brain function, we investigated how within- and between-network functional connectivity were associated with age (**Figure 7**) and overall psychopathology (**Figure 8**). We evaluated functional connectivity matrices that included 400 cortical regions (Schaefer et al., 2018), each assigned to one of the canonical 7 Yeo-Kieren networks (Yeo et al., 2011). Similar to the findings for structural data, age-related variations in network-level connectivity converged to similar patterns following structural and functional QC and data harmonization (**Figure 7B** compared to **Figure 7A**). Following QC and harmonization, we observed an overall increase in within-network connectivity during development for all resting-state networks. However, the amount of age-related increase in within-network connectivity varied between networks with the ventral attention network demonstrating the largest increase in connectivity during development (**Figure 7B**; Partial *R*^2^ = 0.067, *p*_FDR_ < 0.0001). In contrast, between-network functional connectivity decreased with age in most networks. However, these age effects were heterogeneous (**Figure 7B**). For example, connectivity between ventral attention and default mode networks significantly decreased with age (**Figure 7B**; Partial *R*^2^ = −0.065, *p*_FDR_ < 0.0001), while connectivity between ventral attention and dorsal attention networks increased with development (**Figure 7B**; Partial *R*^2^ = 0.046, *p*_FDR_ < 0.0001).

**Figure 7.**
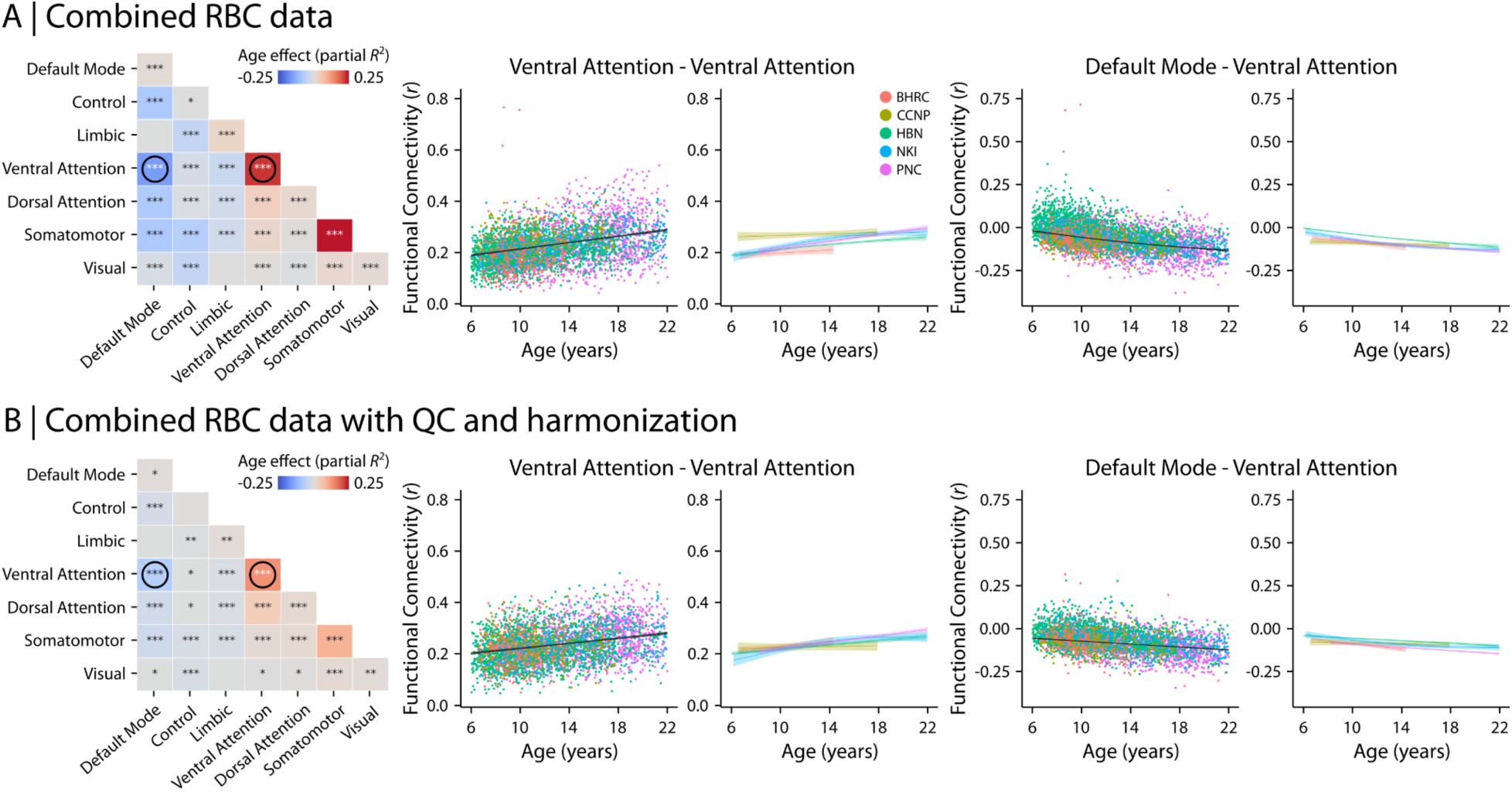
Functional data derivatives are associated with age in youth. To assess the association between brain function and participant age, we examined within- and between-network functional connectivity during development using Generalized Additive Models (GAMs). (A) Aggregated functional data without QC implementation or neuroimaging data harmonization were used to model age-related variations in network-level functional connectivity. The results are depicted in a matrix (left), where the diagonal values correspond to within-network age-effects (partial *R*^2^) and off-diagonal values correspond to between-network age-effects. Asterisks indicate statistical significance of the findings after correcting for multiple comparisons (FDR-corrected *p*-values: * indicates 0.001<*p*<0.05; ** indicates 0.0001<*p*<0.001; *** indicates p<0.0001). Overall, we found an increase in within-network functional connectivity with development (i.e., increased age) while between-network age-effects were more heterogeneous. Example results (marked via circles in the matrix) are shown for within-network connectivity in the ventral attention network (increasing connectivity with development) and between-network connectivity between the default mode and ventral attention networks (decreasing connectivity with development). Study-specific model fits varied between studies, especially for within-network connectivity. (B) We repeated the analyses following QC and imaging data harmonization. Overall, within- and between-network age-effects displayed similar associations as before (with variations in effect size and significance). However, study-specific model fits converged and displayed consistent patterns across studies.

**Figure 8.**
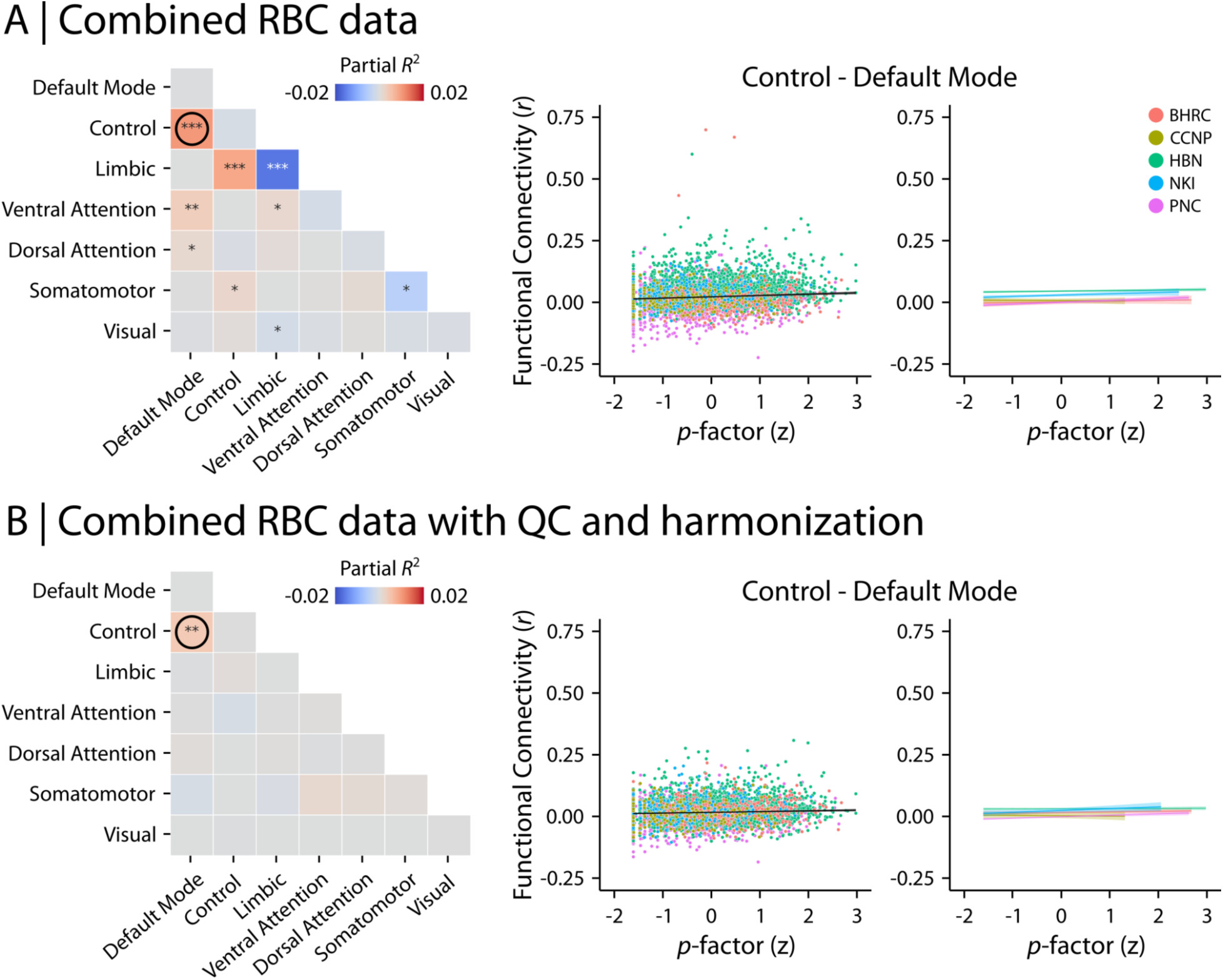
Functional data derivatives are associated with psychopathology in youth. We examined the relationship between within- and between-network functional connectivity and psychopathology (i.e., *p*-factor) using Generalized Additive Models (GAMs). (A) GAM analysis with aggregated functional data without QC implementation or neuroimaging data harmonization identified significance associations between network-level connectivity and *p*-factor in multiple networks. The results are depicted in a matrix (left), where the diagonal values correspond to within-network age-effects (partial *R*^2^) and off-diagonal values correspond to between-network age-effects. Asterisks indicate statistical significance of the findings after correcting for multiple comparisons (FDR-corrected *p*-values: * indicates 0.001<*p*<0.05; ** indicates 0.0001<*p*<0.001; *** indicates p<0.0001). An example relationship (marked via a circle in the matrix) between psychopathology and between-network connectivity is shown for the default model and control (i.e., fronto-parietal) networks. (B) Following QC and imaging data harmonization, we only identified a significant positive association between *p*-factor and between-network connectivity in the default model and control networks, indicating a trend-like increase in connectivity between those two networks with increased psychopathology. Similar to the findings with structural data derivatives (**Figure 6**), these results highlight the importance of rigorous QC and data harmonization protocols in multi-site data analysis.

Finally, we assessed the relationship between *p*-factor and network-level functional connectivity (**Figure 8**). Without QC or harmonization, we found significant associations between *p*-factor and within- and between-network functional connectivity in multiple networks (**Figure 8A**). However, following QC and harmonization, we observed significantly increased connectivity only between default mode and frontoparietal networks with increasing psychopathology (**Figure 8B**; Partial *R*^2^ = 0.004, *p*_FDR_ = 0.0002). As for the features derived from sMRI, we repeated the analyses after implementing QC but before harmonizing imaging data (**Figure S10A** for age effects and **Figure S10B** for psychopathology). Together, these analyses showcase how RBC data can be used and highlight the importance of rigorous QC and data harmonization.

## Discussion

Developmental and psychiatric neuroimaging research often faces significant obstacles: limited sample sizes, variability in image acquisition and psychiatric phenotyping, and inconsistent data analysis workflows (Paus, 2010; Milham, 2012; Poldrack & Poline, 2015; Gilmore et al., 2018; Laird, 2021; Marek et al., 2022). These collectively hinder the generalizability of results, and act as a brake on scientific progress. RBC addresses these challenges by integrating and harmonizing data from over 6,000 participants across five major neurodevelopmental cohorts. We employed advanced harmonization techniques to overcome variability in psychiatric phenotyping and imaging protocols, ensuring that data from different studies can be meaningfully combined with confidence. Furthermore, neuroimaging data with uniform processing are complemented by standardized quality control measures to ensure rigor. Initial results defining consistent patterns of brain development underscore the degree to which RBC facilitates robust and generalizable studies of the developing brain.

RBC is a response to the proliferation of large-scale studies of brain development (Biswal et al., 2010; Somerville et al., 2018; Howell et al., 2019; Satterthwaite et al., 2014; Alexander et al., 2017; Tobe et al., 2022; Liu et al., 2021; Fan et al., 2023). RBC builds on previous data aggregation and harmonization efforts such as the International Neuroimaging Data-sharing Initiative (INDI), the Autism Brain Imaging Data Exchange (ABIDE) Preprocessed, and the ADHD-200 Preprocessed (Biswal et al., 2010; Milham et al., 2012; Di Martino et al., 2013; Craddock et al., 2013a; Bellec et al., 2017), while distinguishing itself through a strong emphasis on harmonization, reproducibility, and data quality. However, this emphasis posed unique and sometimes unforeseen challenges; RBC reflects six years of sustained methodological efforts. Below, we detail what was required as well as lessons learned across several key domains.

### Concise, harmonized psychiatric phenotyping

Harmonizing psychiatric phenotypes remains a major challenge for the field; RBC was no exception. The harmonized dimensions of psychopathology released with RBC were only possible after a series of detailed methodological studies, which used item response theory to harmonize constructs and evaluated a broad range of bifactor models (Hoffmann et al., 2022–2024). The resulting factor scores provide a parsimonious summary of mental health data, disambiguating general and specific dimensions of psychopathology (Reise, 2012; Lahey et al., 2012; Caspi et al., 2014). This approach aligns with the dimensional and hierarchical characterization of psychopathology emphasized in the Hierarchical Taxonomy of Psychopathology (HiTOP; Kotov et al., 2017; Kotov et al., 2019). It should be allowed that recent studies have raised important concerns regarding the alignment of bifactor models of psychopathology with established theoretical frameworks (Watts et al., 2024). Despite potential shortcomings, the validity of the general factor of psychopathology (i.e., *p*-factor) derived from these models is supported by extensive evidence (Caspi & Moffitt, 2018; Sallis et al., 2019; Allegrini et al., 2020; Waszczuk et al., 2021; Hoffmann et al., 2022; Caspi et al, 2024). Importantly, bifactor models also offer pragmatic advantages to researchers, such as parsing inter-item covariance into orthogonal scores (i.e., latent dimensions) that can be used simultaneously in hypothesis testing. When dealing with the inevitable comorbidity of psychopathology symptoms, extracting a general factor to explain them is often the most practical solution. Here we exploited bifactor modeling’s strength in facilitating cross-study comparisons despite the different sampling strategies of the component studies (Hoffmann et al., 2024). The benefit of harmonization was quite evident: overall psychopathology (i.e., *p*- factor) was higher in HBN compared to PNC only in the harmonized data.

### Reproducible image curation

The BIDS format has been a boon for the field (Gorgolewski et al., 2017; Gilmore et al., 2018; Poldrack et al., 2024). However, many studies – like several included in RBC – have not previously been released in BIDS. Furthermore, BIDS meta-data may be inaccurate or missing. This is a critical and under-recognized problem as widely-used image processing pipelines (e.g., “BIDS-apps”) automatically configure workflows based on imaging metadata – leading to processed data that may be inaccurate (e.g., reproducible but wrong). Typically, metadata is corrected manually in an ad-hoc fashion, compromising reproducibility before image processing even begins. This was an unanticipated challenge in RBC – large datasets that required significant meta-data curation. To address this, we created CuBIDS (Covitz et al., 2022), which allows for reproducible BIDS curation. Notably, the detailed summary of metadata provided by CuBIDS also revealed more significant protocol variation in each of the component studies than originally anticipated. Such protocol variation is usually unacknowledged; studies typically report only the intended protocol. Moving forward, the practical consequences of such variation merit evaluation and likely suggest the need for multi-level harmonization methods that can address variation in image acquisition both within and across studies (Beer et al., 2020).

### Uniform image processing and reproducible workflows

In RBC we followed the example of prior data aggregations – such as the ABIDE and ADHD-200 Preprocessed (Craddock et al., 2013a; Bellec et al., 2017) – and released fully processed data. We started by leveraging containerized, open-source pipelines—such as FreeSurfer (Fischl, 2012), sMRIPrep (Esteban et al., 2024), and C-PAC (Craddock et al., 2013b)—to ensure uniform processing of structural and functional MRI data. The need for such uniform processing was underscored after our initial benchmarking study revealed that even seemingly innocuous analytic choices (e.g., template version) can introduce significant variability (Li et al., 2024). Both that study and prior work revealed that the use of global signal regression (GSR) has a very substantial impact on derived features; the impact of GSR on findings remains one of the most common and time-consuming questions addressed in peer review (Macey et al., 2004; Murphy et al., 2009; Saad et al., 2012; Satterthwaite et al., 2013; Power et al., 2015; Murphy & Fox, 2017; Uddin, 2017). We took advantage of the exceptional configurability of C-PAC to execute two high-performance denoising pipelines both with and without GSR. Similarly, C-PAC allowed us to include multiple parcellation systems and quantify connectivity using different measures. Such rich data will facilitate additional methodological studies and will allow investigators to understand how their results may be impacted by important image processing choices without having to re-process the data themselves.

The use of containerized processing pipelines like C-PAC is a major asset for reproducibility – but does not on its own allow for full audit trail. In RBC, we adopted the “FAIRly-big” workflow (Wagner et al., 2022): a framework that uses DataLad (Halchenko et al., 2021) to ensure reproducible processing of large datasets. This framework aligns with the FAIR principles (Findability, Accessibility, Interoperability, and Reusability; Wilkinson et al., 2016) by enabling open, modular, and reusable workflows. However, while we successfully applied the FAIRly-big workflow in RBC, it was not straightforward: it had a steep learning curve despite our team’s significant technical expertise. This experience led our team to develop dedicated software – the BIDS App Bootstrap (BABS; Zhao et al., 2024) – that automates the application of FAIRly-big. While we did not use BABS software for this initial release of RBC, moving forward we anticipate that it will significantly lower the barriers to researchers adopting FAIRly-big and DataLad in their own work.

### Quality control

One critical yet often-overlooked factor in cross-study reproducibility is the role of quality control (QC) and variation in QC-based sample selection procedures. Even when identical processing pipelines and QC metrics are used, differences in inclusion and exclusion criteria can result in different samples included in data analysis, potentially leading to divergent findings from the same dataset. The ambiguity of QC selection criteria creates opportunities for accidental or deliberate p-hacking, further complicating cross-study comparisons. To address these challenges, we provide extensive harmonized QC metrics and recommended QC guidelines for using RBC data. These include both categorical QC measures that can be used for sample selection as well as continuous measures of data quality that can be used as model covariates.

While RBC provides harmonized QC metrics and standardized guidelines, our analyses highlight significant remaining challenges. For example, we observed that participants who are younger or had higher levels of psychopathology tended to have poorer data quality. This raises important concerns about how QC procedures may inadvertently reduce generalizability to populations of the most interest. Ultimately, while our QC efforts confirm many findings from prior studies (e.g., Alexander et al., 2017; Tobe et al., 2022), they also underscore the trade-offs inherent in balancing data quality with inclusivity. Moving forward, improvements in image acquisition – such as online motion correction with vNavs (Tisdall et al., 2016) – will be essential for ensuring that studies can effectively capture populations of the greatest interest and need.

### Convergent developmental findings and ongoing challenges in translational psychiatry

As an initial evaluation of the utility of RBC, we examined the associations of structural and functional imaging features with age. Aligning with a large body of prior work (Sowell et al., 2003; Sowell et al., 2004; Shaw et al., 2008; Raznahan et al., 2011; Wierenga et al., 2014; Xu et al., 2015; Bethlehem et al., 2022), we found evidence for a decline in CT, SA, and GMV in all datasets following careful QC and harmonization. Consistent with a rich literature from lifespan network neuroscience (Kelly et al., 2008; Supekar et al., 2009; Fair et al., 2009; Zuo et al., 2010; Vogel et al., 2010; Hagmann et al., 2010; Hagmann et al., 2012; Collin & van den Heuvel, 2013; Menon, 2013; Betzel et al., 2014; Cao et al., 2014; Di Martino et al., 2014; Luna et al., 2015; Luo et al., 2024), we also found evidence for a decline in between-network functional connectivity paired with increases in within-network connectivity. These findings suggest that resting-state functional networks become more segregated and specialized during development (Fair et al., 2007; Betzel et al., 2014; Gu et al., 2015; Luna et al., 2015; Keller et al., 2023; Luo et al., 2024). Notably, these results were far less consistent prior to harmonization and QC, emphasizing how variation in data quality and acquisition parameters may obscure even robust effects of brain development.

In contrast to the highly consistent developmental findings, the existing literature on associations between major dimensions of psychopathology and imaging features have been more varied (Goodkind et al., 2015; Snyder et al., 2017; Barch, 2017; Kaczkurkin et al., 2019; Mewton et al., 2021; Mattoni et al., 2021; Romer et al., 2021; Parkes et al., 2021; Royer et al., 2024). These inconsistencies may be due to small sample sizes, limited reliability, and biological heterogeniety (Yarkoni, 2009; Poldrack et al., 2017; Zuo et al., 2019; Milham et al., 2021; Marek et al., 2022; Gell et al., 2024). Echoing at least part of the existing literature (Goodkind et al., 2015; Romer et al., 2017; Snyder et al., 2017; Kaczkurkin et al., 2019; Mewton et al., 2021; Royer et al., 2024), we found that reduced SA and GMV were linked to higher overall psychopathology. Additionally, we found that higher overall psychopathology was associated with greater connectivity between the default mode and frontoparietal networks, suggesting a loss of normative developmental network segregation (Kelly et al., 2008; Di Martino et al., 2014; Satterthwaite et al., 2015; Xia et al., 2017; Bassett et al., 2018; Baker et al., 2019; Kebets et al., 2019; Royer et al., 2024). These results are not surprising in that they are consistent with multiple published reports– this consistency and the very large sample used butresses confidence in our results. However, in contrast to the larger effects of brain development, it should be emphasized that the effect sizes of associations with psychopathology were small. Such small effect sizes remain a major challenge for the field, highlighting the need for more sensitive imaging measures, better methods for characterizing psychopathology, and approaches for parsing heterogeneity in links between brain and behavior (Woo et al., 2017; Zuo et al., 2019; Feczko et al., 2019; Sui et al., 2020; Kaczkurkin et al., 2020; Milham et al., 2021; Tian & Zalesky, 2021; Gell et al., 2024).

Notably, our analyses of brain development and links to psychopathology also highlighted the importance of QC and data harmonization. For example, our initial analysis of combined RBC data without QC or neuroimaging data harmonization identified significant associations between CT and psychopathology in multiple cortical regions. However, those associations were no longer significant following QC and data harmonization. We observed similar effects in the fMRI data. These findings underscore that large samples are necessary but not sufficient – high quality, harmonized data is also required.

### Limitations and future directions

There are several methodological and technical limitations that must be considered when using RBC. First, RBC provides extensive demographics and phenotypic data (e.g., *p*-factor) that can be accessed and used without any restrictions. However, there are other behavioral, clinical, and demographic variables that are not released as part of the RBC dataset due to specific data privacy and DUA protocols set by the component studies. Although such phenotypic measures are not included in RBC, this information can be requested directly from each study. Second, the majority of the data included in RBC are cross-sectional datasets, with the exception of a smaller subset of individuals who have follow-up longitudinal data. Future work is required to provide public large longitudinal datasets to systematically study neurodevelopmental trajectories of brain and behavior data within the same individuals over time. For example, the Adolescent Brain Cognitive Development (ABCD; Volkow et al., 2018) study provides a powerful resource, tracking a large cohort from childhood through adolescence. Notably, ABCD data can only be released via NIH-approved methods that authenticate access; as such it cannot be included in the RBC sharing model. Third, while RBC provides a relatively diverse data resource and incorporates data from five prominent neurodevelopmental datasets spanning three different continents, it is not fully representative of diverse populations. Neuroimaging studies often lack adequate inclusion of minority groups, highlighting the need for future data collection and sharing efforts to build larger and more diverse samples that can be used to reliably capture neurodevelopmental patterns at the population level. Fourth, parent- or caregiver-reports were the primary source of psychopathology measures in RBC. While using parent-reported measures of psychopathology provides valuable insights and is considered a reasonable approach, these reports reflect only one perspective and often diverge from youth self-reports (Xavier et al., 2022; Jones et al., 2024). Future research should prioritize developing measures with higher parent-youth concordance or relying on direct youth-reports for more accurate and reliable assessments. Finally, RBC’s neuroimaging data consist of structural and functional MRI data. Future efforts are required to provide large-scale neurodevelopmental datasets that include multiple neuroimaging modalities, such as diffusion-weighted MRI (DWI) studies of brain microstructure and arterial spin-labeled (ASL) MRI measures of cerebral perfusion.

Moving forward, RBC’s utility will be amplified by its adherence to FAIR principles – Findability, Accessibility, Interoperability, and Reusability. RBC data is easily ***Findable*** and readily ***Accessible***: all raw and fully processed RBC data are publicly shared via the International Neuroimaging Data-sharing Initiative (INDI) and are accessible without any DUA requirements. By openly releasing de-identified data and removing barriers associated with cumbersome DUAs, RBC accelerates scientific discovery. RBC data ***Interoperability*** is ensured through the use of standard data structures like BIDS and tidy tabular derivative data along with well-documented, open-source imaging pipelines. Moreover, RBC data can be redistributed without restriction, ensuring that it is ***Reusable***. For example, researchers can use RBC data to develop tools or integrate RBC data with other datasets. Crucially, such efforts can be shared alongside RBC on INDI. RBC is also accompanied by a version-controlled website to help facilitate data access and maintenance (https://reprobrainchart.github.io/). We provide a “quick-start guide” on how to access and download RBC data on the RBC website (https://reprobrainchart.github.io/docs/get_data). We also provide detailed analysis workflows that allow users to replicate the results reported here (see “Data and Code availability”). For support, researchers can post questions to INCF NeuroStars using the RBC tag (https://neurostars.org/tag/rbc). Taken together, RBC accelerates large-scale, robust, and reproducible research in developmental and psychiatric neuroscience.

## Methods

### Study description

The RBC project contains data from 5 large studies of brain development (age range: 5-85 years old) from *N*=6,346 participants (*N*=2,869 Female). All studies included structural and functional Magnetic Resonance Imaging (MRI) data as well as phenotypic data. Studies were conducted in Brazil, China, and the United States of America. Specific studies include: Brazilian High Risk Cohort (BHRC; *n*=610; Salum et al., 2014), Developmental Chinese Color Nest Project (CCNP; *n*=195; Liu et al., 2021; Fan et al., 2023), Healthy Brain Network (HBN; *n*=2,611; Alexander et al., 2017), Nathan Kline Institute–Rockland Sample (NKI; *n*=1,329; Tobe et al., 2022), and Philadelphia Neurodevelopmental Cohort (PNC; *n*=1,601; Satterthwaite et al., 2014; Satterthwaite et al., 2016).

BHRC (Salum et al., 2014) is a sample of children and adolescents attending school in Brazil (Porto Alegre and São Paulo cities) that is aimed to be a random sample of the state-funded school-based community in addition to children with increased family risk of mental disorders. The study was approved by the ethics committee of the University of São Paulo.

CCNP (Liu et al., 2021; Fan et al., 2023) is a sample of children and adolescents that is aimed to represent the population residing in multiple cities of China with varying economies and from different regions of the country (https://ccnp.scidb.cn/en). The ethical approval for this study was obtained from the Institutional Review Board of the Chinese Academy of Sciences (CAS) Institute of Psychology and Beijing Normal University.

HBN (Alexander et al., 2017) is a sample of children and adolescents residing in the New York City area (United States of America) that is aimed to represent a diverse sample of healthy and help-seeking individuals with heterogeneous metrics of developmental psychopathology. The HBN study included data from four different acquisition sites: Staten Island (SI - Mobile Scanner), Rutgers University (RU), The City University of New York (CUNY), and Citigroup Biomedical Imaging Center (CBIC). The study was approved by the Chesapeake Institutional Review Board.

NKI–Rockland Sample (Tobe et al., 2022) is aimed to represent a lifespan sample of individuals with varying demographic distributions residing in the United States. The Institutional Review Board approved this project at the Nathan Kline Institute.

PNC (Satterthwaite et al., 2014; Satterthwaite et al., 2016) is a community sample of children and adolescents residing in the greater Philadelphia area (United States) that is aimed to represent a diverse developmental sample. The study was approved by the Institutional Review Boards of the University of Pennsylvania and the Children’s Hospital of Philadelphia.

Written informed consents were obtained from all participants (or their parents or legal guardians) by each study separately as part of the study-specific data collection procedure.

T1-weighted structural MRI and resting-state, task, and movie-watching functional MRI data were included in the RBC dataset. All scans were defaced and de-identified to ensure ethical compliance and protect participants’ privacy. Data acquisition parameters varied by study and data acquisition sites. Detailed information about structural and functional MRI data acquisitions are summarized in **Tables S4–5**.

### Phenotypic data harmonization

RBC includes psychiatric phenotyping data for each study, which were assessed using one of two different phenotypic questionnaires. The Child Behavior Checklist (CBCL; Achenbach & Rescorla, 2001) was used in the BHRC, CCNP, HBN, and NKI studies, while the GOASSESS interview (Calkins et al., 2015) was used in the PNC study. To ensure that measures of psychopathology were consistent across study, we used a bifactor modeling strategy to harmonize differences between samples that used the same instrument (i.e., the CBCL) as well as differences between disparate instruments (i.e., GOASSESS vs. CBCL). The CBCL is a 120- item parent-report assessment of emotional and behavioral symptoms over the past 6 months, answered on a 3-point scale (0 = not true, 1 = somewhat/sometimes true, and 2 = very true/often). It encompasses eight syndromes: anxious-depressed, withdrawn-depressed, somatic complaints, rule-breaking behavior, aggressive behavior, social problems, thought problems, and attention problems (Achenbach & Rescorla, 2001). To harmonize CBCL with GOASSESS, CBCL scores of 1 and 2 were collapsed to generate a binary-scaled variable compatible with GOASSESS (i.e., 0 or 1). The GOASSESS is a structured screening interview administered to collateral informants (usually a caregiver) by trained assessors. It contains 112 unconditioned screening items based on DSM-IV constructs, including symptoms of mood disorders (Major Depressive Episode, Manic Episode), anxiety disorders (Generalized Anxiety Disorder, Separation Anxiety Disorder, Specific Phobia, Social Phobia, Panic Disorder, Agoraphobia, Obsessive-Compulsive Disorder, Post-traumatic Stress Disorder), Attention Deficit/Hyperactivity Disorder (ADHD), behavioral (Oppositional Defiant Disorder, Conduct Disorders) and eating disorders (Anorexia, Bulimia), and suicidal thinking and behavior. Items are scored as 0 (absent) or 1 (ever present). The instrument is abbreviated and modified from the epidemiologic version of the NIMH Genetic Epidemiology Research Branch Kiddie-SADS, and its development is described and tested elsewhere (Calkins et al., 2015).

We tested a sequence of harmonization procedures to minimize between-cohort and between-questionnaire differences across studies, as well as to disentangle general and specific aspects of psychopathology. In a previous work, we thoroughly tested the impact of different bifactor model configurations on the resulting factor scores (Hoffmann et al., 2022) in addition to six item-matching strategies that best harmonize different questionnaires (Hoffmann et al., 2024). We then tested the best bifactor model configuration using several parameters to identify the model that best harmonized CBCL and GOASSESS questionnaires used in RBC in another previous work (Hoffmann et al., 2023). In brief, 12 CBCL bifactor models were first identified from previous literature, which varied in item and factor configurations (Hoffmann et al., 2022). The impact of these modeling choices was small for the general psychopathology factor (i.e., *p*- factor), but quite marked for the specific factors (Hoffmann et al., 2022). We then tested 6 item-matching strategies to harmonize items between questionnaires (i.e., CBCL and GOASSESS), where we found that the expert-based 1-to-1 semantic item-matching performed best for item harmonization (Hoffmann et al., 2024). Finally, the extent to which the CBCL–GOASSESS harmonized models were similar to the original models was assessed across different models (Hoffmann et al., 2023). We selected the McElroy model as it: (1) demonstrated measurement invariance between the questionnaires; (2) retained the majority of original items during the harmonization process; (3) included harmonized items that were endorsed in all samples; and (4) was among the best four harmonized bifactor models in terms of factor reliability and authenticity (i.e., generated factor scores fairly correlated and close to the factor scores from full item set models) (Hoffmann et al., 2022; Hoffmann et al., 2023, Hoffmann et al., 2024).

In our previous RBC harmonization studies and for the publicly released RBC dataset, all factor scores were estimated with confirmatory factor analysis (CFA) using delta parameterization and Weighted Least Squares with diagonal weight matrix with standard errors and mean- and variance-adjusted Chi-square test statistics (WLSMV) estimators. RBC studies were used as clusters in the CFA in the bifactor model that included all RBC data. Analysis were carried out in Mplus 8.6 (Muthén & Muthén, 2017) and implemented in R version 4.0.3 using the MplusAutomation package (Hallquist & Wiley, 2018), which was also used to extract factor scores generated in Mplus using maximum a posteriori method. The global model fit was evaluated using root mean square error of approximation (RMSEA), comparative fit index (CFI), Tucker–Lewis index (TLI), and standardized root mean square residual (SRMR). RMSEA lower than 0.060 and CFI or TLI values higher than 0.950 indicate a good-to-excellent model. SRMR lower than or equal to 0.080 indicates acceptable fit, and lower than 0.060 in combination with previous indices indicates good fit (Hu & Bentler, 1999). Only individuals that filled the CBCL and GOASSESS on the day of neuroimaging were included in the publicly released RBC dataset.

### Reproducible neuroimaging data curation and workflows

We curated all raw neuroimaging data and metadata in a fully-reproducible fashion that conforms with the Brain Imaging Data Structure (BIDS; Gorgolewski et al., 2016) using the Curation of BIDS (CuBIDS; Covitz et al., 2022) software package. CuBIDS provides a workflow for identifying unique combinations of imaging data acquisition parameters based on metadata, summarizing the heterogeneity in an MRI BIDS dataset, and reproducibly modifying scan filenames to reflect information about their imaging parameters. These curation steps are critical given that preprocessing pipelines designed for BIDS datasets (e.g., “BIDS Apps”; Gorgolewski et al., 2017) automatically configure workflows based on a dataset’s accompanying metadata. However, potential inaccuracies or missing information in metadata may lead to incorrectly configured pipelines that will run without any errors (i.e., pipelines that are reproducible but wrong). CuBIDS facilitates the identification of such inaccuracies or missing information in metadata and categorizes imaging data in subgroups (i.e., unique parameter groups) based on the heterogeneity of their metadata.

We applied CuBIDS to RBC data and generated a CuBIDS summary data file for each study included in the RBC dataset. The CuBIDS summary tables are provided in **Data S1–5**. CuBIDS was then used to rename each BIDS data file based on the scanning parameters of the identified parameter groups, such that the new file names indicated the source of the variance between different parameter groups. Notably, all data curation steps were fully tracked via CuBIDS’s wrapped use of DataLad (Halchenko et al., 2021) throughout, ensuring the tracking of version history and yielding a complete audit trail. After categorizing data based on heterogeneity of their acquisition parameters, we used an automated procedure implemented in CuBIDS to create an example dataset with one subject from each acquisition group for each study. The example dataset was then used to closely examine the heterogeneity of data and make necessary adjustments to the data curation step (e.g., if there were errors or missing information in metadata). Eventually, the example dataset was used to test image processing pipelines to ensure that they performed well on each combination of acquisition parameters present in the dataset. Finally, to run the RBC neuroimaging data through modality specific preprocessing pipelines, we employed the FAIRly-big workflow (Wagner et al., 2022). FAIRly-big is a DataLad-based open source framework that is suitable for reproducible processing of large-scale datasets. We adapted the FAIRly-big workflow for RBC image processing to ensure that all data preparation and processing were accompanied by a full audit trail in DataLad (Halchenko et al., 2021).

### Structural MRI data processing and quality control

Structural MRI (sMRI) data were processed using FreeSurfer v6.0.1 (Fischl, 2012) and sMRIPrep v0.7.1 (Esteban et al., 2024), yielding commonly used measures of brain structure. Specifically, structural images underwent correction for intensity non-uniformity and skull-stripping with ANT’s brain extraction workflow. Brain surfaces were then reconstructed using FreeSurfer. RBC provides full FreeSurfer outputs as well as tabulated data parcellated using 35 anatomical, functional, and multimodal atlases that were included to align with the functional image processing. Atlases include the Desikan Killiany (Desikan et al., 2006), Glasser (Glasser et al., 2016a), Gordon (Gordon et al., 2016), and multiple resolutions of the Schaefer (Schaefer et al., 2018) parcellation among others. Specific features include commonly used measures of brain structure such as regional surface area, cortical thickness, gray matter volume, and folding and curvature indices. Moreover, summary brain measures such as total intracranial volume, ventricle size, and mean and standard deviation of various measures (e.g., cortical thickness, surface area) are provided for the whole brain and per hemisphere. Tabulated data are also accompanied by .json files describing each structural feature in detail.

An important feature of RBC is the emphasis on harmonized measures of neuroimaging data quality control (QC). To achieve consistent quality ratings across all studies in RBC, every structural image was manually evaluated by 2-5 expert raters using Swipes for Science, a web application for binary image classification (Keshavan et al., 2019). The expert rating workflow consisted of seven phases (**Figure 3**). Overall, 4 two-dimensional slices per participant (two axial and two sagittal slices extracted per structural scan) were created to rate each scan. To ensure consistent slice selection across participants, the anatomical images were registered (linear rigid body) to the MNI152 template space prior to slice selection. Expert raters (experience in brain imaging: mean = 5.63 years; SD = 5.76 years) evaluated a total of 28,780 slices and assigned “Pass” or “Fail” to those slices based on visual inspection only. A “Pass” rating would be given if an image was deemed of sufficient quality for skullstriping and segmentation of cerebrospinal fluid (CSF), white, and gray matter. Slices were presented to the raters in random order.

More specifically, Phase 1 involved generating the ground truth for 200 images to either “Pass” (rating of 1) or “Fail” (rating of 0). Two senior raters evaluated these images and agreed on 96% of the slices. The images with disagreement were further evaluated after skulltripping and brain segmentation, using Configurable Pipeline for the Analysis of Connectomes (C-PAC; Craddock et al., 2013b) pipeline. The expert raters then reached 100% concordance through consensus of either “Pass” or “Fail” for each slice. In Phase 2, these ground-truth annotated images (i.e., outcome of Phase 1) were used to train four raters until they reached at least 85% concordance with the ground-truth. In Phase 3, all raters manually rated 10% of images and the reliability of ratings was assessed across raters. In Phase 4, the raters were divided into two groups with balanced experience and evaluated 20% of the remaining images independently, such that no overlapping images were assessed by the two groups. In Phase 5, the reliability of ratings were re-assessed (similar to Phase 3), where all raters manually rated another 10% of the remaining images. In Phase 6, the raters were divided into two groups again to rate 20% of the remaining images (similar to Phase 4). Finally, in Phase 7, a new set of images that were not included in the first 6 phases were evaluated by raters. To avoid sequence type or scanner parameters biases, each Swipes for Science instance for each phase contained data from all data acquisition sites and studies.

Following manual expert ratings, an overall QC determination score of “Pass”, “Artifact”, or “Fail” was assigned to each structural image as the final scan quality based on the average rating across raters. “Pass” or “Fail” labels were assigned to images if all raters were in agreement about their quality. For example, if all raters agreed that a given image was of adequate quality (i.e., all raters assigned “Pass” or rating of 1 to the image), the average rating across raters was equal to 1 and the image was labeled as “Pass”. Similarly, if all raters assigned “Fail” or rating of 0 to an image, the image was labeled as “Fail” (i.e., average rating of 0 across raters). If there was any disagreement between raters about a given image’s quality (i.e., a combination of 1 and 0 ratings for the image), the “Artifact” label was assigned to that image (i.e., an average value between 0 and 1). Summary information on the number of participants with structural scans before and after applying RBC’s recommended QC is available in **Figures S1–S5**.

In addition to the manual ratings, we calculated the Euler number from FreeSurfer (Rosen et al., 2018). The Euler number is calculated as part of the FreeSurfer processing pipeline and reflects the topological complexity of the initial reconstructed cortical surface (Rosen et al., 2018). Euler number is calculated as the sum of the vertices and faces subtracted by the number of faces. Lower values below an ideal value of 2 indicate more defects in surface reconstruction. Euler number can be used as a continuous measure of structural MRI QC (e.g., can be included as a covariate in downstream analysis) whereas manual ratings provide categorical QC labels. Previous reports have demonstrated that the Euler number is an accurate, fully-automated measure of data quality, capturing variation in data quality within the coarse categories provided by manual ratings (Rosen et al., 2018; Klapwijk et al., 2019; Elyounssi et al., 2022; Elyounssi et al., 2023; Tobe et al., 2022).

### Functional MRI data processing and quality control

As noted above, preprocessing steps were carried out in DataLad (Halchenko et al., 2021) using the FAIRly-big workflow (Wagner et al., 2022) to track provenance and ensure the re-executability of processing workflows for all RBC studies. Following guidelines from extensive benchmarking and harmonization studies (Li et al., 2021), functional MRI (fMRI) data were preprocessed using Configurable Pipeline for the Analysis of Connectomes (C-PAC v1.8.5.dev1; Craddock et al., 2013b), with a pipeline configuration file specifically developed for RBC (C-PAC Developers, 2022) including a fixed random state to maximize reproducibility. This configuration produces commonly used measures of functional MRI data such as fully-processed fMRI time-series and functional connectivity matrices (e.g., Pearson and partial correlations between processed regional time-series). In addition, the functional data derivatives include regional measures based on processed time-series such as Regional Homogeneity (ReHo; Zang et al., 2004), Amplitude of Low Frequency Fluctuation (ALFF; Zang et al., 2007), and fractional ALFF (fALFF; Zou et al., 2008). Processed time-series and functional connectivity are available in parcellated format using 17 different atlases, including AAL (Tzourio-Mazoyer et al., 2002), Glasser (Glasser et al., 2016a) and Schaefer (Schaefer et al., 2018) parcellations. ReHo, ALFF, and fALFF are available in volumetric MNI space (MNI152NLin6ASym).

For anatomical preprocessing, the configuration specified C-PAC’s reimplementation of the NiWorkflows ANTs brain extraction workflow (NiPreps Developers, 2021; Esteban et al., 2020; Tustison et al., 2021) and segmentation using FSL-FAST (Zhang et al., 2001) with a 0.95 threshold for each tissue type. Anatomical registration was configured to use antsRegistration (Tustison et al., 2021) on skull-stripped images at 1mm isotropic resolution with parameters specified in the configuration file (C-PAC Developers, 2022).

Functional registration was configured to first use C-PAC’s reimplementation of the NiWorkflows reference image estimation method (Center for Reproducible Neuroscience, 2023a; Esteban et al., 2021) with boundary-based registration performed on skull-stripped images to MNI152NLin6ASym-template space using FSL-FLIRT (Jenkinson & Smith, 2001; Jenkinson et al., 2002) with binarized partial volume white matter masks. It then used C-PAC’s reimplementation of the fMRIPrep single-step resampling workflow (fMRIPrep developers, 2020; Esteban et al., 2019) at 2mm isotropic resolution. Motion statistics were calculated before slice-timing correction using the previously established reference image and FSL-MCFLIRT (Jenkinson et al., 2002) for motion estimation. Where field maps were present, distortion correction was performed using FSL-FUGUE (Jenkinson et al., 2012) or FSL-TOPUP (Jenkinson et al., 2012) depending on the type of field maps. Functional masking was configured to use a reference image from TemplateFlow (Ciric et al., 2022) and C-PAC’s reimplementation of the NiWorkflows BOLD masking method (Center for Reproducible Neuroscience, 2023b; Esteban et al., 2021). A mean functional image was also generated using AFNI 3dTstat (Cox, 1996). The first 2 timepoints were excluded from analysis.

AFNI 3dDespike (Cox, 1996) was run on template-space images. Prior to nuisance regression, the brain mask, CSF mask, and white matter mask were eroded using C-PAC’s reimplementation of the NiWorkflows utility interface for converting tissue probability masks into regions of interest (Center for Reproducible Neuroscience, 2023c; Esteban et al., 2021). Two nuisance regression strategies (named “36-parameter” and “aCompCor”, with global signal regression (GSR) or aCompCor as part of the strategy, respectively and exclusively) were run in parallel in template space (the full specific nuisance regression configurations can be found in the configuration file, C-PAC Developers, 2022).

Time-series extraction (C-PAC developers, 2023a) was performed and correlation matrices were generated using Nilearn’s “correlation” and “partial correlation” methods (nilearn developers, 2018; Nilearn contributors, 2024) separately for each of the specified region of interest atlases. ALFF (Zang et al., 2007) and fALFF (Zou et al., 2008) were calculated in template space using a sequence of AFNI tools (C-PAC developers, 2023b, 2023c; Cox, 1996). Regional Homogeneity (Zang et al., 2004) was calculated in template space using NiBabel (Brett et al., 2019) and NumPy (Harris et al., 2020) in Python (C-PAC developers, 2023d).

C-PAC outputs also include extensive measures of quality control, such as various in-scanner motion parameters (e.g., framewise displacement), functional image to structural image (e.g., to T1-weighted scan or MNI template) registration and normalization quality parameters, generated for each preprocessed BOLD image using C-PAC’s reimplementation (C-PAC developers, 2023e) of the xcpEngine quality control dictionary (Ciric et al., 2019). As part of the RBC data release, we performed an initial quality assurance procedure using measures of in-scanner motion quantified as Framewise Displacement (FD) and normalization quality quantified as cross correlation from normalization of functional image to template image. Functional MRI runs with a median FD less than or equal to 0.2 and normalized cross correlation greater than or equal to 0.8 were considered of adequate quality. These thresholds were selected after extensive manual review of images. Similar to structural data, the majority of functional scans passed RBC’s QC thresholds. Summary information on the number of participants with functional scans before and after applying RBC’s recommended QC is available in **Figures S1– S5**. Overall, about 90% of RBC data passed both structural and functional QC guidelines.

Following the initial structural and functional data QC procedure, three different versions of the RBC dataset were publicly released and are accessible depending on the user’s choice of QC threshold: (1) structural images with “Pass” label only and functional images that passed the motion and image normalization QC threshold; (2) structural images with “Pass” and “Artifact” labels and functional images that passed the motion and image normalization QC threshold; (3) all structural and functional images including QC failures. Versions (1) and (2) are recommended by RBC; however, users may use Version (3) for research on QC or to apply their desired QC procedures using the extensive structural and functional QC information accompanying RBC data.

### Example analysis workflow with RBC data

In addition to providing a publicly available data resource, we illustrated the utility of RBC data and the increased statistical power of the aggregated sample. Specifically, we examined the relationship between derived structural and functional neuroimaging features and both participant age and general psychopathology. Furthermore, we evaluated whether those associations are influenced by RBC’s QC protocols and imaging data harmonization.

#### Neuroimaging data harmonization

RBC provides both raw and processed neuroimaging data for each dataset. However, due to differences in scanners and sequences used in data collection between studies, there is significant technical variance between studies and sites that requires harmonization. Therefore, users may want to statistically harmonize neuroimaging data across studies and data acquisition sites. However, neuroimaging data harmonization remains an ongoing research topic in the field with a growing number of statistical methods that aim to effectively harmonize data between multiple sites and studies. Given the lack of consensus on a single neuroimaging data harmonization technique, we released unharmonized neuroimaging data in RBC. This allows researchers to implement their method of choosing to harmonize the imaging data based on their specific research questions. In addition, given that statistical harmonization models require specification of covariates that are hypothesis-dependent, researchers can define the covariates they want to include in the harmonization analysis. While RBC’s released neuroimaging data is unharmonized, below we provide an example workflow to harmonize the imaging data and illustrate its utility in our analysis. This workflow can be tailored by RBC users for their specific hypotheses.

We used CovBat-GAM (Johnson et al., 2007; Fortin et al., 2017; Fortin et al., 2018; Pomponio et al., 2020; Chen et al., 2022) to harmonize structural and functional MRI data across data acquisition sites. Specifically, we used the “covfam” function with Generalized Additive Model (GAM) from ComBatFamility in R 4.2.2. The “covfam” function uses CovBat (Correcting Covariance Batch Effects; Chen et al., 2022) to harmonize the mean and covariance of data across multiple batches (e.g., sites, datasets). In our developmental analysis, data acquisition sites were treated as batches. Covariates included in harmonization were age as a smooth term (to account for linear and nonlinear age effects), and sex and data quality (i.e., Euler number for structural data and median FD for functional data) as linear terms. We used a separate harmonization for the psychopathology analyses, where general psychopathology from the bifactor model (*p*-factor) was added as a linear term to the model. CovBat-GAM yields harmonized structural and functional data while protecting effects of model covariates. To evaluate the effect of harmonization, we repeated analysis before and after data harmonization.

#### Generalized Additive Models (GAM)

We used Generalized Additive Models (GAMs) to delineate linear and nonlinear developmental effects (i.e., age effects) and assess the relationship between neuroimaging data derivatives and general psychopathology (i.e., *p*-factor) (Pomponio et al., 2020; Sydnor et al., 2023; Luo et al, 2024). The analysis was performed using the “mgcv” package in R 4.2.2. To ensure all studies contributed to the analysis, a subset of aggregated RBC data within the age range of 6- 22 years old was used for this analysis. Structural features included cortical thickness (CT), surface area (SA), and gray matter volume (GMV), and functional features included between- and within-network resting-state functional connectivity. Structural and functional data parcellated into 400 brain regions using the Schaefer-400 atlas (Schaefer et al., 2018). Functional networks were defined using the 7 Yeo-Krienen networks for the parcellated functional data (Yeo et al., 2011). Within-network connectivity was estimated as the average functional connectivity between regions from a given network and all the other regions from the same network. Between-network connectivity was estimated as the average connectivity between regions from a given network and regions from all the other networks. Note that we used the baseline functional scans of individuals who were scanned at multiple time points (e.g., BHRC and NKI). For individuals with multiple functional data at the baseline, we used the average functional connectivity across scans at the baseline.

Separate GAMs were fit for functional and structural features as dependent variables. Each GAM included age as a smooth term and sex and data quality as linear covariates. The linear covariate for data quality was the Euler number for structural data and median FD for functional data. For models evaluating associations with overall psychopathology, the *p*-factor was also included in GAMs as a linear term. In all GAM analyses, the maximum basis complexity was set to 3 for the smooth term (i.e., age) to avoid overfitting. The code invocation used to fit the model was: ‘mgcv::gam(feature ∼ s(age, k=3, fx=F) + factor(sex) + data_quality)’. The GAM formula also included *p*-factor as a linear term for psychopathology analysis. The Restricted Maximum Likelihood (REML) approach was used to estimate the smoothing parameters in GAMs. We performed the analyses at two resolutions: (1) whole-brain models, where we examined average structural and functional features across the cortex for each participant; (2) regional and network level GAMs, where we evaluated regional structural features and network-level functional features (e.g., mean within- or between-network connectivity) for each participant.

The effect size of associations between imaging features and age were estimated as partial *R*^2^. We quantified partial *R*^2^ as the difference in Residual Sum of Squares (SSE) between the full model that included the smooth term for age and a reduced model with no age term (i.e., only including model covariates) normalized by SSE of the reduced model. Normalization by SSE of the reduced model highlights the relative contribution of the predictor (e.g., age) to the reduced model. The linear effect size for the relationship between *p*-factor and neuroimaging features was also estimated in a similar manner, where partial *R*^2^ was defined as the difference in SSE between the full model and a reduced model without *p*-factor. We used a signed version of partial *R*^2^, such that the sign reflected the directionality of observed effects (increase or decrease in neuroimaging features with increasing age or *p*-factor). To obtain the directionality of partial *R*^2^ for age analysis, we calculated the mean derivative of the model fit for the smooth term (i.e., age). A positive sign was assigned to partial *R*^2^ if the mean derivative was positive, reflecting an overall increasing trend between the neuroimaging feature of interest and age, whereas a negative sign was assigned to partial *R*^2^ where the mean derivative was negative. To obtain the directionality of partial *R*^2^ for *p*-factor analysis, we used the sign of the *t*-value associated with the linear *p*-factor term. The statistical significance of the associations between neuroimaging features and age or *p*-factor was assessed using analysis of variance (ANOVA) between the full model and reduced model that excluded the term of interest. Results were corrected for multiple comparisons by controlling for the false discovery rate (FDR correction; *Q*<0.05).

To assess how neuroimaging data QC and harmonization impacted our findings, we repeated all structural and functional GAM analyses with three different versions of data: First, we used all neuroimaging data without excluding individuals based on QC thresholds and without applying neuroimaging data harmonization. Second, we used structural MRI data from individuals with “Pass” or “Artifact” structural QC (i.e., excluding scans with “Failed” structural QC) and functional data that passed functional QC thresholds (median FD < 0.2 and normalization cross correlation > 0.8). However, we did not apply neuroimaging data harmonization in this version. Third and finally, we used neuroimaging data that passed structural and functional QC (as above) and performed GAM analyses after applying neuroimaging data harmonization.

### Data sharing protocol and code availability

All harmonized phenotypes as well as raw and processed structural and functional images are openly shared via DataLad and the International Neuroimaging Data-sharing Initiative (INDI). The released RBC data are accessible without any Data User Agreement (DUA) requirements and can be downloaded using DataLad via https://github.com/ReproBrainChart. Moreover, all processing pipelines are shared using Docker containers for frictionless portability across platforms (https://github.com/ReproBrainChart) along with the analysis workflows and data used in the present study (https://github.com/ReproBrainChart/rbc-analysis-template). RBC data release is also accompanied by a website to help facilitate data access and maintenance (https://reprobrainchart.github.io/). The RBC website provides additional information and guidelines on how to access the data. Follow-up queries on RBC are closely monitored on INCF Neurostars (https://neurostars.org/tag/rbc).

## Acknowledgments

RBC was funded by the National Institute of Mental Health (NIMH) via grant R01MH120482. Additional support was provided by R37MH125829, R01EB022573, R01MH112847, R01MH113550, RF1MH121867, R01MH123550, U24NS130411, P50MH109429, R01MH123440, K08MH079364. GS was supported by a postdoctoral fellowship from the Canadian Institutes of Health Research (CIHR). We would like to thank the AWS Open Data Sponsorship Program for storage support. CCNP receives funding support from the STI 2030–the major projects of the Brain Science and Brain-Inspired Intelligence Technology (2021ZD0200500), the National Basic Science Data Center “Interdisciplinary Brain Database for In-vivo Population Imaging” (ID-BRAIN), the Key-Area Research and Development Program of Guangdong Province (2019B030335001), the Start-up Funds for Leading Talents at Beijing Normal University, the Beijing Municipal Science and Technology Commission (Z161100002616023, Z181100001518003), the Major Project of National Social Science Foundation of China (20&ZD296), the CAS-NWO Programme (153111KYSB20160020), the Guangxi BaGui Scholarship (201621), the National Basic Research (973) Program (2015CB351702), the Major Fund for International Collaboration of National Natural Science Foundation of China (81220108014), the Chinese Academy of Sciences Key Research Program (KSZD-EW-TZ-002) and the National Basic Research Program (2015CB351702). The Philadelphia Neurodevelopmental Cohort was supported by NIMH RC2 MH089983 and RC2 MH089924. BHRC was funded by Conselho Nacional de Desenvolvimento Científico e Tecnológico (CNPq grant numbers 573974/2008-0 and 465550/2014-2), Fundação de Amparo à Pesquisa do Estado de São Paulo (FAPESP grant numbers: 2008/57896-8, 2013/08531-5, 2014/50917-0, 2020/06172-1, 2021/05332-8, 2021/12901-9) to the National Institute of Development Psychiatric for Children and Adolescent (INPD), the European Research Council under the European Union’s Seventh Framework Programme (FP7/2007-2013)/ERC grant agreement no 337673.

