## Supplementary figures and tables for "Reproducible Brain Charts: An open data resource for mapping brain development and its associations with mental health"

**Affiliations:**
[1] Penn Lifespan Informatics and Neuroimaging Center, University of Pennsylvania, Philadelphia, PA, USA
[2] Department of Psychiatry, Perelman School of Medicine, University of Pennsylvania, Philadelphia, PA, USA
[3] Lifespan Brain Institute (LiBI) of Penn Medicine and CHOP, University of Pennsylvania, Philadelphia, PA, USA
[4] Child Mind Institute, New York, NY, USA
[5] Department of Neuropsychiatry, Universidade Federal de Santa Maria (UFSM), Santa Maria, Brazil
[6] Graduate Program in Psychiatry and Behavioral Sciences, Universidade Federal do Rio Grande do Sul (UFRGS), Porto Alegre, Brazil
[7] National Institute of Developmental Psychiatry & National Center for Innovation and Research in Mental Health, Brazil
[8] Care Policy and Evaluation Centre, London School of Economics and Political Science, London, UK
[9] Department of Public Health Sciences, Medical University of South Carolina, Charleston, SC, USA
[10] Department of Bioengineering, Stanford University, Stanford, CA, USA
[11] Department of Neuroscience, Columbia University, New York, NY, USA
[12] Khoury College of Computer Sciences, Northeastern University, Boston, MA, USA
[13] Center for Biomedical Imaging and Neuromodulation, Nathan Kline Institute, Orangeburg, NY, USA
[14] Department of Psychiatry, NYU Grossman School of Medicine, New York, NY, USA
[15] Department of Radiology, University of Pennsylvania, Philadelphia, PA, USA.
[16] Center for Biomedical Image Computing and Analytics, University of Pennsylvania, Philadelphia, PA, USA
[17] Department of Psychiatry, Federal University of São Paulo (UNIFESP), São Paulo, Brazil
[18] Department of Psychology, University of Washington, Seattle, WA, USA
[19] eScience Institute, University of Washington, Seattle, WA
[20] Penn Statistics in Imaging and Visualization Center, Department of Biostatistics, Epidemiology, and Informatics, University of Pennsylvania, Philadelphia, PA, USA
[21] Department of Psychology, Columbia University, New York, NY, USA
[22] Developmental Population Neuroscience Research Center, IDG/McGovern Institute for Brain Research, Beijing Normal University, Beijing, China
[23] ADHD Outpatient Program & Developmental Psychiatry Program, Hospital de Clinicas de Porto Alegre, Universidade Federal do Rio Grande do Sul (UFRGS), Porto Alegre, Brazil
[24] Medical Council UNIFAJ & UNIMAX, Brazil

* Equal contribution

### Supplementary Figures:

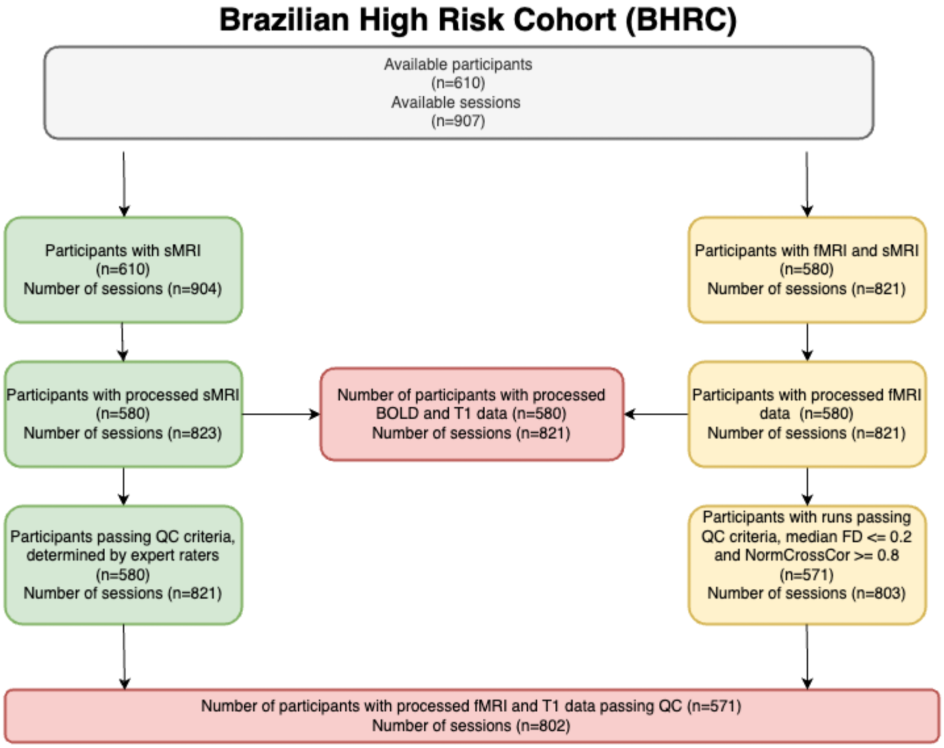


**Figure S1. Quality control workflow for Brazilian High Risk Cohort (BHRC) dataset |** Majority of the neuroimaging data in BHRC pass RBC's quality control guidelines (see Table S6 for more details).

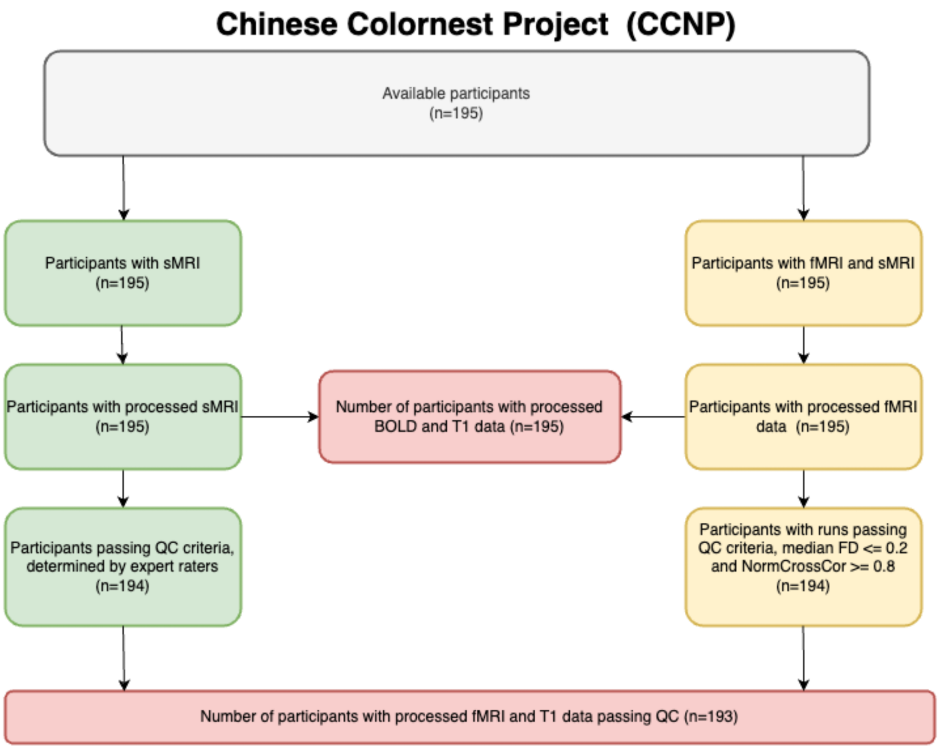


**Figure S2. Quality control workflow for Developmental Chinese Colornest Project (CCNP) dataset |** Majority of the neuroimaging data in CCNP pass RBC's quality control guidelines (see Table S6 for more details).

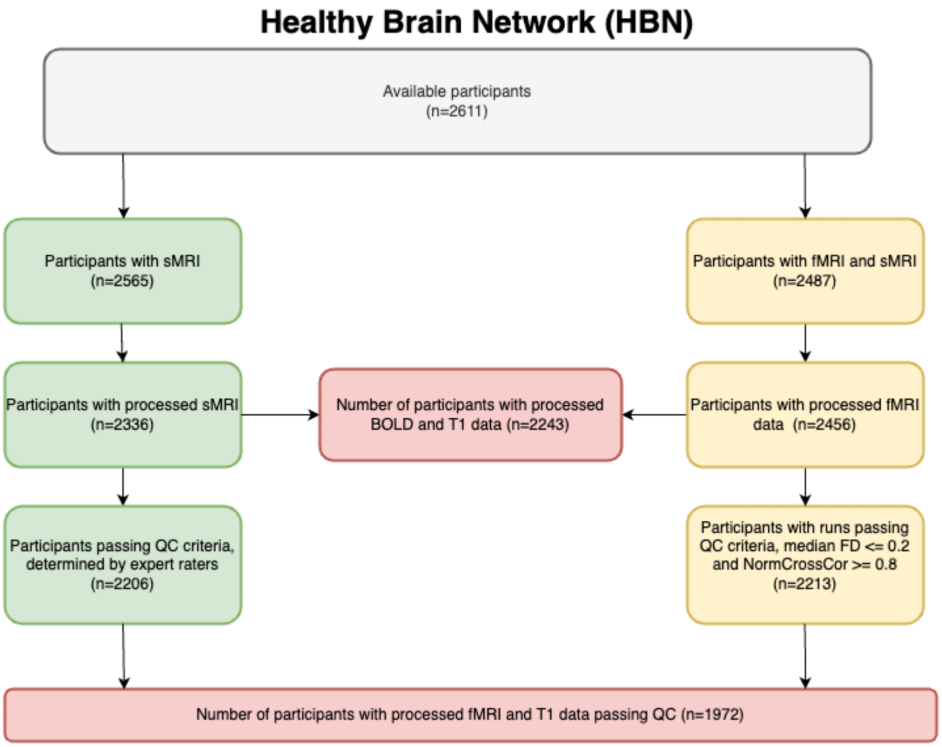


**Figure S3. Quality control workflow for Healthy Brain Network (HBN) dataset |** Majority of the neuroimaging data in HBN pass RBC's quality control guidelines (see Table S6 for more details).

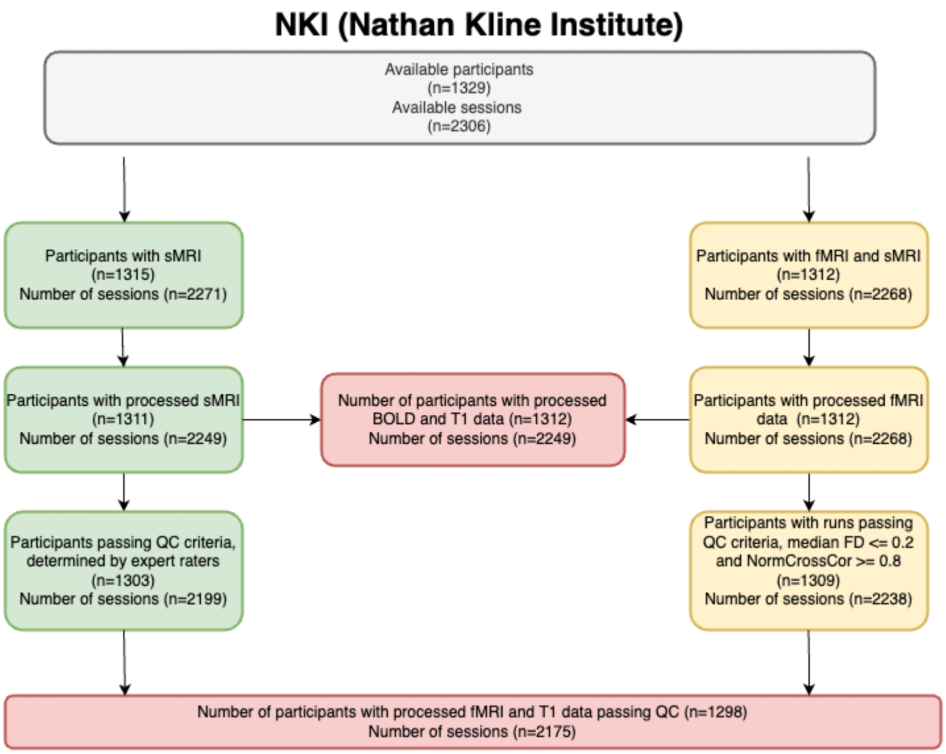


**Figure S4. Quality control workflow for Nathan Kline Institute – Rockland Sample (NKI) dataset |** Majority of the neuroimaging data in NKI pass RBC's quality control guidelines (see Table S6 for more details).

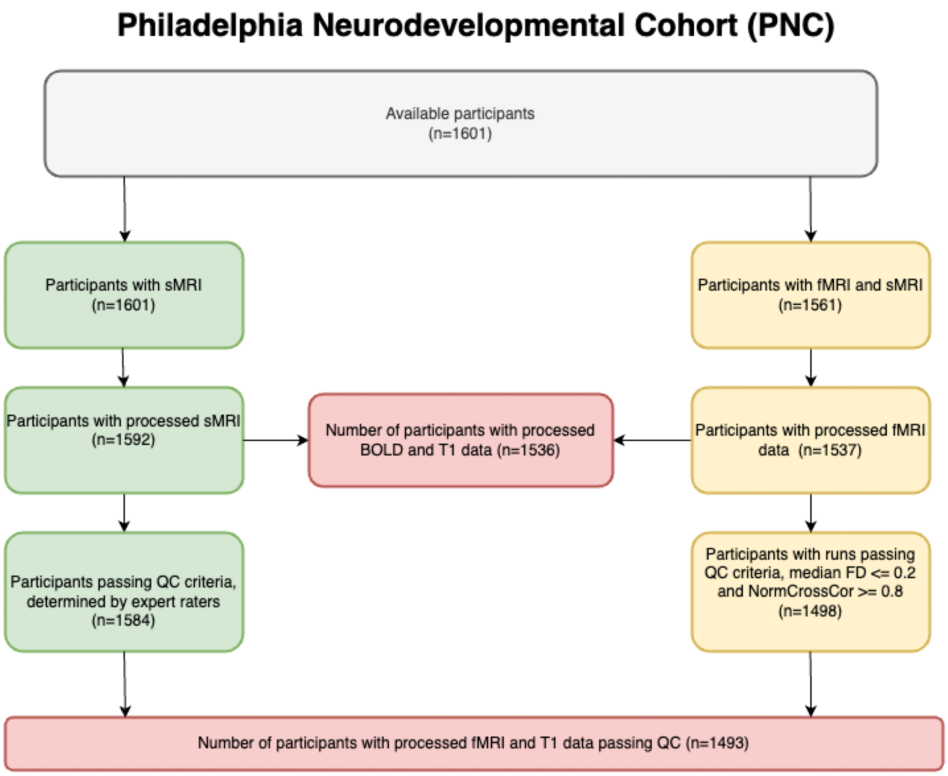


**Figure S5. Quality control workflow for Philadelphia Neurodevelopmental Cohort (PNC) dataset |** Majority of the neuroimaging data in PNC pass RBC's quality control guidelines (see Table S6 for more details).

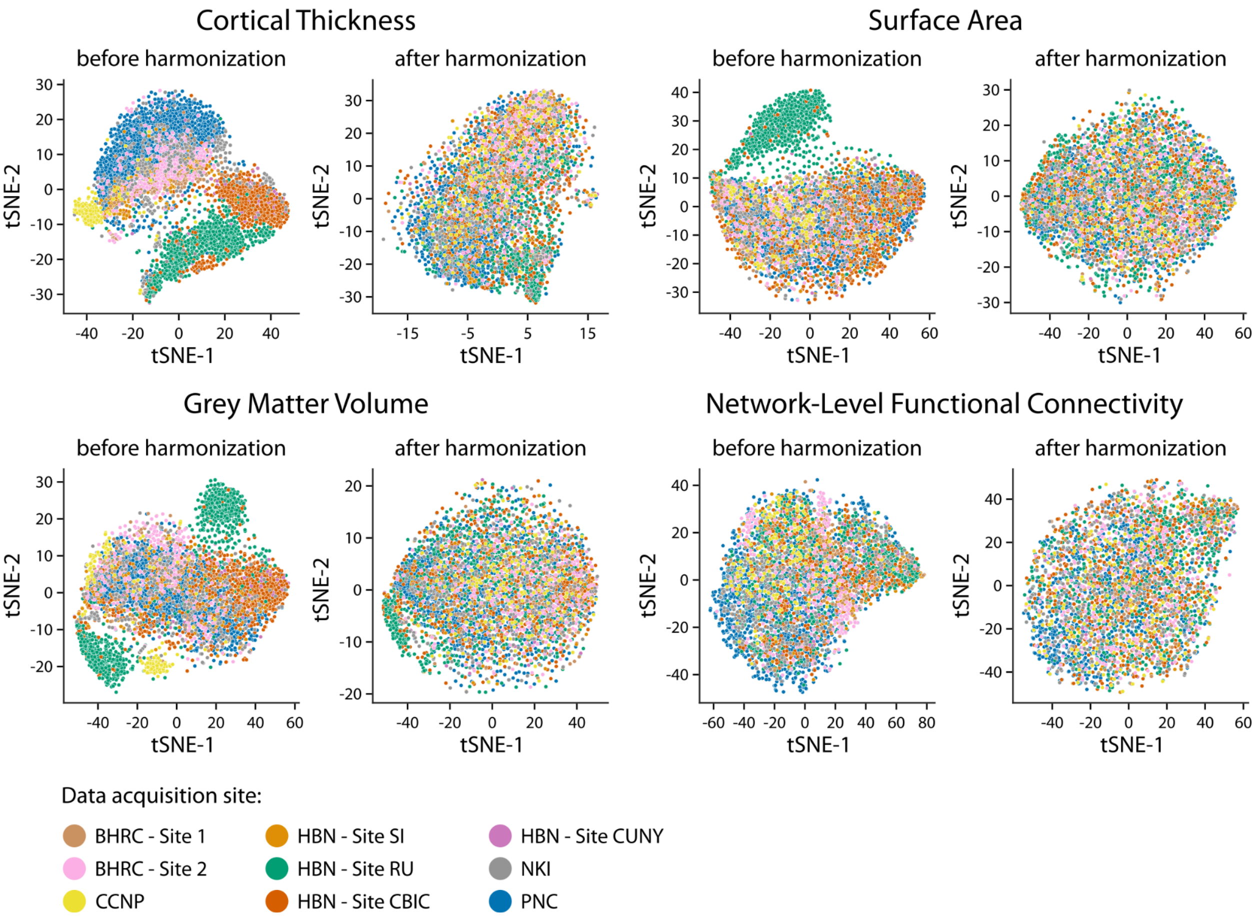


**Figure S6. Impact of neuroimaging data harmonization using CovBat-GAM |** tSNE scatter plots are used to illustrate the impact of CovBat-GAM (Johnson et al., 2007; Fortin et al., 2017; Fortin et al., 2018; Pomponio et al., 2020; Chen et al., 2022) on harmonizing structural and functional MRI data across acquisition sites while preserving model covariate effects (e.g., age, sex, data quality, psychopathology). The plots depict neuroimaging features—structural (cortical thickness, surface area, grey matter volume) and functional (network-level functional connectivity)—aer color-coded by acquisition site in a low dimensional space. After harmonization, site-specific clusters observed before harmonization largely disappear, indicating a more homogeneous distribution of data points.

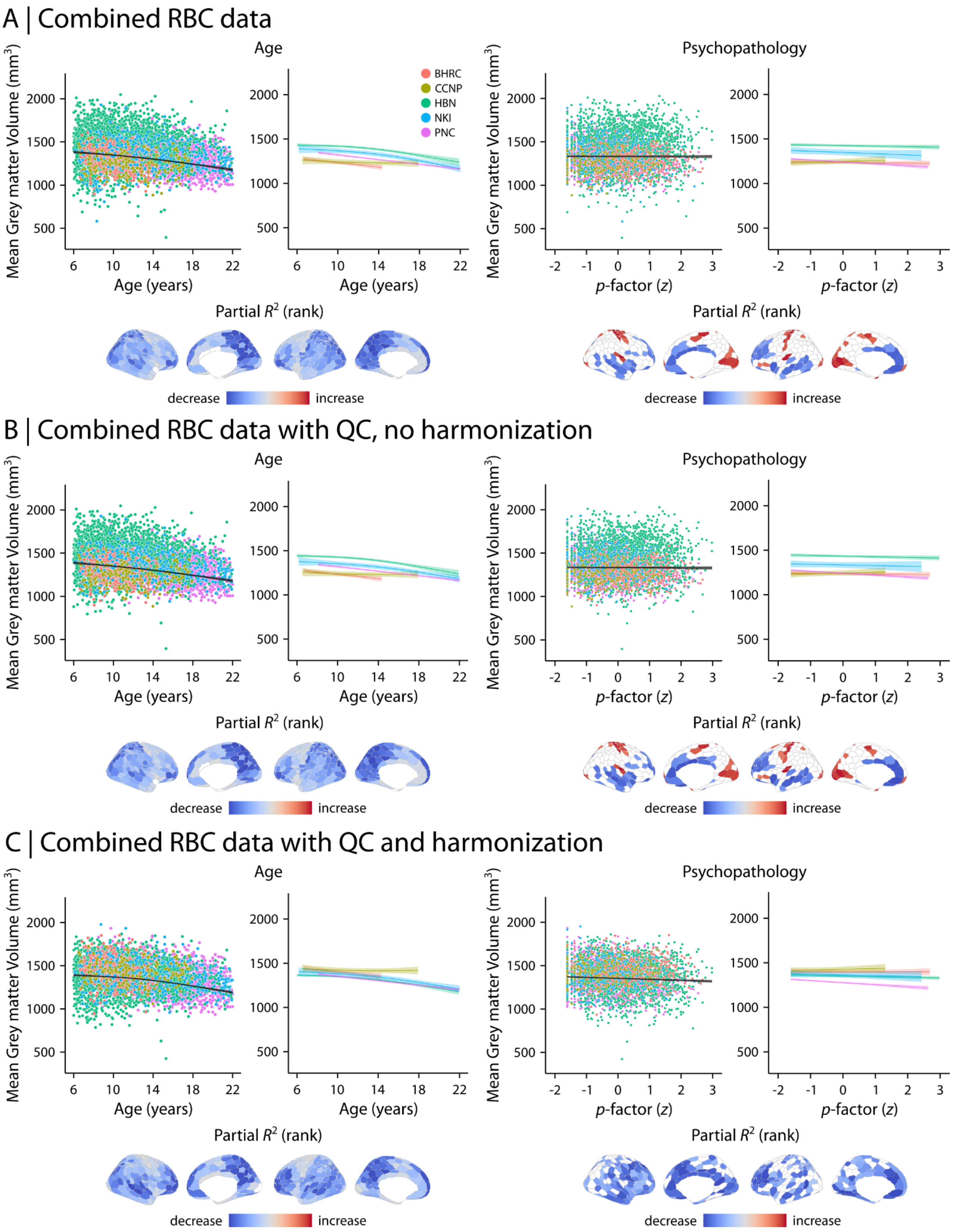


**Figure S7. Grey matter volume predicts age and psychopathology in youth |** Generalized Additive Models (GAMs) were used to examine how grey matter volume (GMV) is associated with participant age (left column) and psychopathology (i.e., *p*-factor; right column) in RBC. (A)

We first examined  aggregated RBC data without QC or neuroimaging data harmonization. We found an overall decrease in mean GMV during development (with increasing age) while mean GMV remained mostly unchanged with respect to psychopathology. However, whole-brain associations between GMV and age as well as psychopathology varied markedly between studies as demonstrated by study-specific model fits (depicted with color coded curves). We also examined regional effects using separate models for each brain region. Regional effects were quantified using ranked partial $R^2$ and are depicted on the cortical surface after correcting for multiple comparisons (FDR-corrected). These regional analyses demonstrated that GMV decreases in most brain regions with age, while it increases in the sensorimotor cortex.  We found overall psychopathology was associated with reduced GMV in prefrontal cortices. (B) To assess the effects of QC, we first repeated the analysis after excluding data with “Fail” structural QC determination score, but without harmonizing neuroimaging data. (C) We then repeated the analysis after harmonizing data using CovBat-GAM in addition to excluding data with “Fail” structural QC determination score to assess the effects of both QC and data harmonization. Following QC and data harmonization, the developmental effects on mean GMV as well as the association between psychopathology and GMV became more similar between studies. Results reveal an overall decrease in mean GMV with development. Regional analyses identified heterogeneous developmental effects on the cortex. GMV decreased in the majority of brain regions with age (more prominent decrease in medial parietal, lateral prefrontal, and temporal regions). Moreover, regional analysis identified significant declines in GMV with increased psychopathology, which were most prominent in medial and lateral prefrontal cortices.

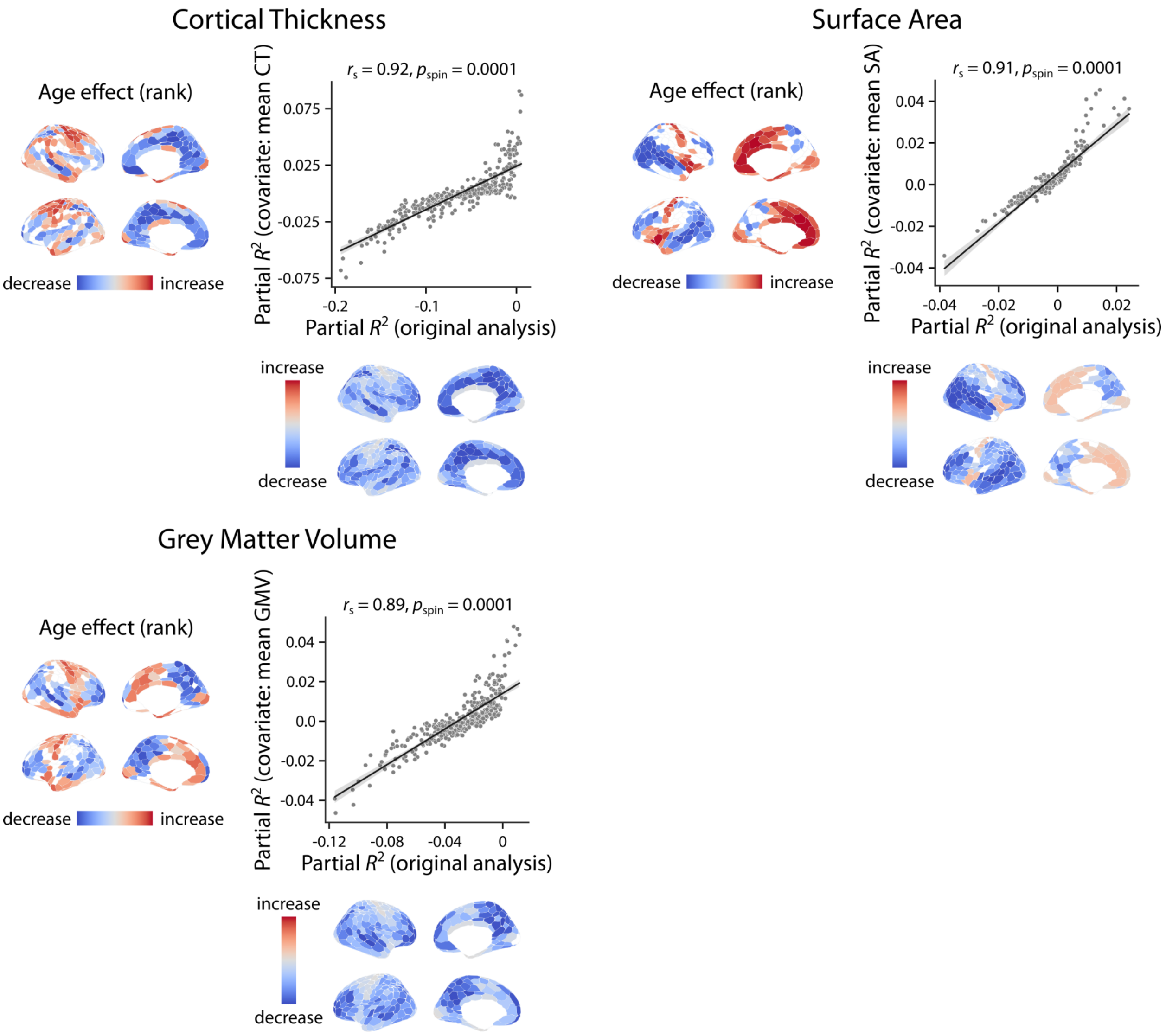


**Figure S8. Development of brain structure after controlling for global effects |** Developmental GAM analysis were repeated while controlling for global effects by including mean CT, SA, and GMV in addition to other covariates (i.e., sex and Euler number). Similar to the original analysis (**Figure 5** and **Figure S7**), regional age effects were quantified using ranked partial $R^2$ and are depicted on the cortical surface after correcting for multiple comparisons (FDR-corrected). The findings demonstrated that regional variations were generally consistent with the original analyses after controlling for global effects. The associations between original age effects and age effects from the sensitivity analysis were quantified using Spearman rank correlation ($r_s$). Statistical significance of the reported associations was assessed using 10,000 spatial autocorrelation- preserving null models (Alexander-Bloch et al., 2018; Markello & Misic., 2021). Linear fit lines were added to the scatter plots for visualization purposes only.

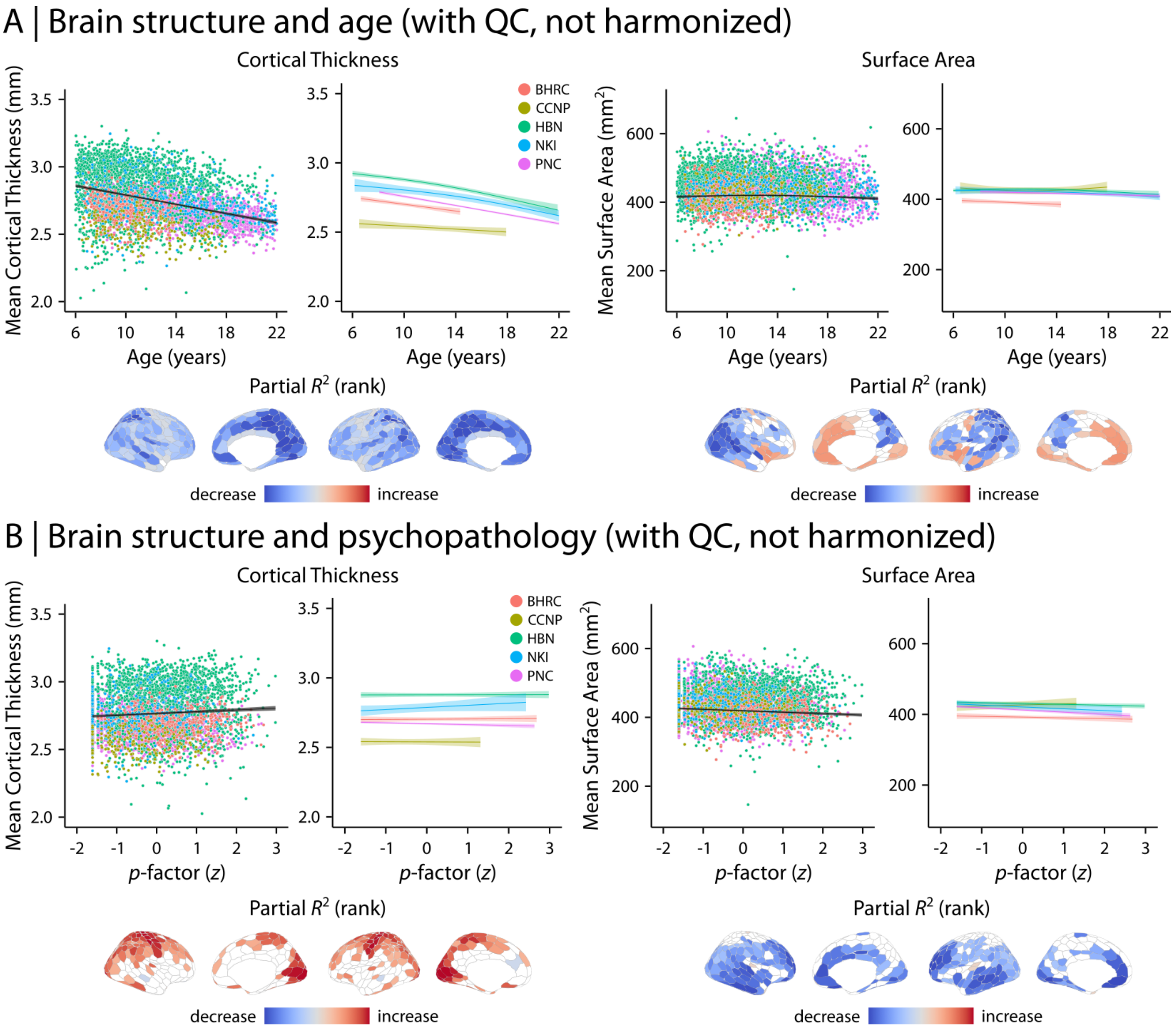


**Figure S9. Impact of quality control on associations between brain structure, age, and psychopathology |** To assess the effects of quality control (QC) on how cortical thickness (CT) and surface area (SA) relate to age (A) and psychopathology (B), we repeated the analysis demonstrated in **Figures 5-6** after excluding data with “Fail” structural QC determination score, but without harmonizing neuroimaging data. Although QC removed unreliable data points from both analyses, the resulting patterns remained similar to the original analysis shown in **Figure 5A** for age and **Figure 6A** for psychopathology, where no QC or data harmonization was implemented. These findings underscore the importance of neuroimaging data harmonization in multi-site datasets in addition to rigorous QC.

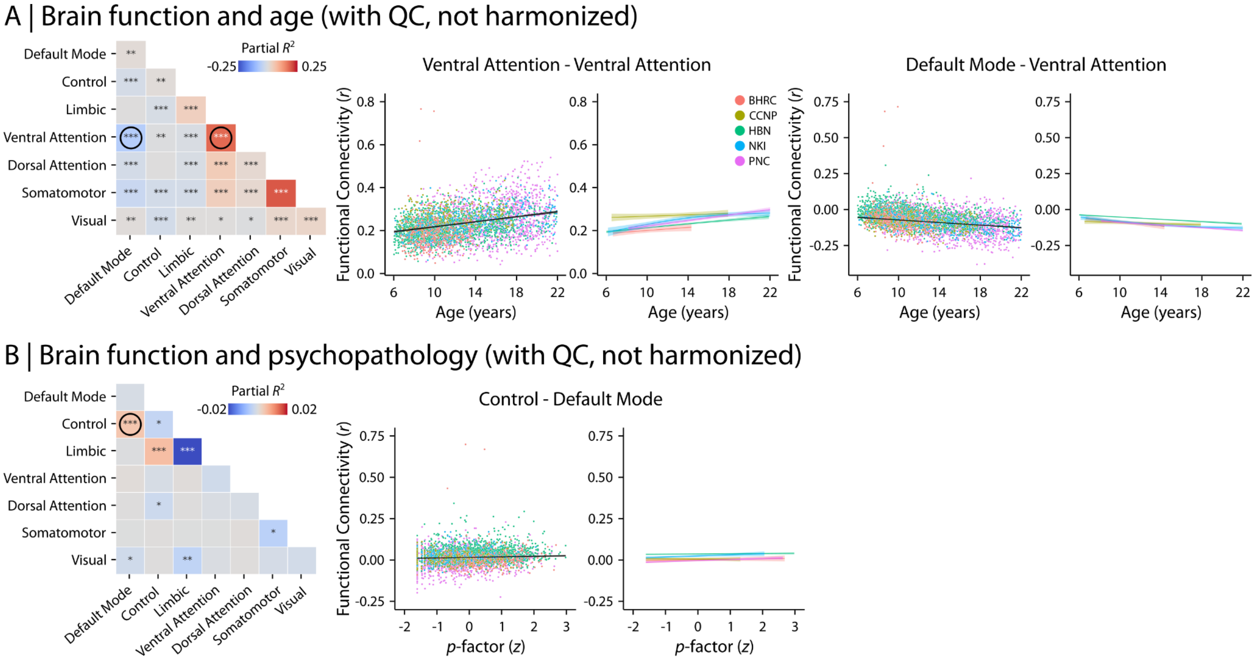


**Figure S10. Impact of quality control on associations between brain function, age, and psychopathology |** To assess the effects of quality control (QC) on how within- and between-network connectivity relate to age (A) and psychopathology (B), we repeated the analysis demonstrated in **Figures 7-8** after excluding data with “Fail” structural and functional QC scores, but without harmonizing neuroimaging data. Although QC removed unreliable data points from both analyses, the resulting patterns remained similar to the original analysis shown in **Figure 7A** for age and **Figure 8A** for psychopathology, where no QC or data harmonization was implemented. These findings underscore the importance of neuroimaging data harmonization in multi-site datasets in addition to rigorous QC.

**Supplementary Tables:**

**Table S1. RBC aggregates five diverse neurodevelopmental datasets**

| Dataset | Number of sites | Sample size | Number of sessions | Age range (mean), years | Structural MRI sessions | Functional MRI sessions |
|---|---|---|---|---|---|---|
| **Brazil High Risk Cohort (BHRC)** | 2 | 610 | 907 | 5.8-14.3 (9.8) | 904 | 821 |
| **Developmental Chinese Color Nest Project (CCNP)** | 1 | 195 | 195 | 6.5-17.9 (11.9) | 195 | 195 |
| **Healthy Brain Network (HBN)** | 4 | 2,611 | 2,611 | 5.0-21.9 (10.4) | 2,565 | 2,487 |
| **Nathan Kline Institute – Rockland Sample (NKI)** | 1 | 1,329 | 2,306 | 6.2-85.6 (36.6) | 2,271 | 2,268 |
| **Philadelphia Neurodevelopmental Cohort (PNC)** | 1 | 1,601 | 1,601 | 8.1-23.1 (14.9) | 1,601 | 1,561 |
| **Total** | | 6,346 | 7,620 | | 7,536 | 7,332 |

**Table S2. CBCL-GOASSESS harmonized bifactor model fit indices |** CBCL, Child Behavior Checklist. BHRC, Brazilian High-Risk Cohort Study for Mental Conditions; HBN, Healthy Brain Network; NKI, Nathan Kline Institute-Rockland Sample; CCNP, developmental component of the Chinese Color Nest Project; PNC, Philadelphia Neurodevelopmental Cohort. RMSEA, Root Mean Square Error of Approximation; CFI, Comparative Fit Index; TLI, Tucker-Lewis Index; SRMR, Standardized Root Mean-square Residual.

| **McElroy model: harmonized items** | **RMSEA** | **RMSEA: 90% CI** | **CFI** | **TLI** | **SRMR** |
|---|---|---|---|---|---|
| **All studies in the model** | 0.011 | [0.009, 0.013] | 0.974 | 0.968 | 0.064 |
| **BHRC study (*n* = 772)** | 0.035 | [0.029, 0.040] | 0.968 | 0.961 | 0.063 |
| **HBN study (*n* = 3252)** | 0.046 | [0.044, 0.048] | 0.953 | 0.943 | 0.065 |
| **NKI study (*n* = 372)** | 0.042 | [0.034, 0.050] | 0.953 | 0.942 | 0.100 |
| **CCNP study (*n* = 181)** | 0.028 | [0.000, 0.044] | 0.964 | 0.956 | 0.144 |
| **PNC study (*n* = 1600)** | 0.057 | [0.053, 0.060] | 0.950 | 0.938 | 0.085 |

**Table S3. CBCL-GOASSESS harmonized McElroy bifactor model using BHRC, HBN, NKI, CCNP and PNC samples |** CBCL, Child Behavior Checklist; BHRC, Brazilian High-Risk Cohort Study for Mental Conditions; HBN, Healthy Brain Network; NKI, Nathan Kline Institute-Rockland Sample; CCNP, developmental component of the Chinese Color Nest Project; PNC, Philadelphia Neurodevelopmental Cohort. PUC, percent of uncontaminated correlations; ECV, explained common variance;  SS, proportion of common variance of the items in each factor which is due to that factor; SG, ECV proportion of common variance of the items in each specific factor which is due to the specific factor; GS, proportion of common variance of the items in each specific factor which is due to the general factor; OmegaH, omega-hierarchical; H, index of construct replicability (> 0.8 suggests a well-defined latent variable); FD, factor determinacy (> 0.9 indicate that the factor score can be used).

| CBCL | | GOASSESS | | Factors and factor loadings | | | |
|---|---|---|---|---|---|---|---|
| Content | Item | Content | Item | P-factor | Attention/ hyperactivity | Internalizing | Externalizing |
| Can't concentrate, can't pay attention for long | CBCL_8 | Have trouble paying attention on activities that you were doing | ADD_011 | 0.537 | 0.840 | | |
| Can't sit still, restless or hyperactive | CBCL_10 | Have difficulty sitting still for more than a few minutes at a time | ADD_020 | 0.607 | 0.494 | | |
| Confused or seems to be in a fog | CBCL_13 | Been told that you did not seem to be listening when they spoke to you | ADD_016 | 0.536 | 0.333 | | |
| Poor school work | CBCL_61 | Make careless mistakes in school work or other activities | ADD_014 | 0.562 | 0.428 | | |
| Cries a lot | CBCL_14 | Cried a lot or felt like crying | DEP_002 | 0.551 | | 0.194 | |
| Fears certain animals, situations, or places, other than sch | CBCL_29 | Afraid of any other things or situations | PHB_008 | 0.388 | | 0.397 | |
| Fears he/she might think or do something bad | CBCL_31 | Bothered by fear that you would do/say something bad without intending to | OCD_004 | 0.515 | | 0.348 | |
| Too fearful or anxious | CBCL_50 | Worry a lot more than most people your age | GAD_002 | 0.504 | | 0.627 | |
| Self-conscious or easily embarrassed | CBCL_71 | Afraid being the center of attention and were concerned something embarrassing might happer | SOC_005 | 0.509 | | 0.353 | |
| Too shy or timid | CBCL_75 | Really shy with people meeting new people going to parties or doing things in front of others | SOC_001 | 0.369 | | 0.332 | |
| Talks about killing self | CBCL_91 | You have thought about killing yourself | SUI_002 | 0.659 | | 0.057 | |
| Unhappy, sad, or depressed | CBCL_103 | You felt sad or depressed most of the time | DEP_001 | 0.713 | | 0.221 | |
| Worries | CBCL_112 | Have been a worrier | GAD_001 | 0.392 | | 0.663 | |
| Cruelty, bullying, or meanness to others | CBCL_16 | Often bully others | CDD_005 | 0.568 | | | 0.635 |
| Lying or cheating | CBCL_43 | Got into trouble with adults like lying or stealing | CDD_001 | 0.591 | | | 0.286 |
| Physically attacks people | CBCL_57 | Try to hurt someone with a weapon | CDD_007 | 0.567 | | | 0.618 |
| Sets fires | CBCL_72 | Set fires break into cars or destroy someone else's property on purpose | CDD_003 | 0.337 | | | 0.245 |
| Stubborn, sullen, or irritable | CBCL_86 | Irritable or grouchy or get angry because you thought that things were unfair | ODD_006 | 0.785 | | | 0.175 |
| Sudden changes in mood or feelings | CBCL_87 | You felt unusually grouchy cranky or irritable | MAN_007 | 0.859 | | | 0.013 |
| Sulks a lot | CBCL_88 | Felt grouchy irritable or in a bad mood most of the time | DEP_004 | 0.790 | | | -0.092 |
| Temper tantrums or hot temper | CBCL_95 | Losing temper arguing with adults or being grouchy or irritable with them | ODD_001 | 0.736 | | | 0.296 |
| Threatens people | CBCL_97 | Threaten someone | CDD_008 | 0.613 | | | 0.562 |
| *Index* | | | | | | | |
| PUC | | | | 0.662 | | | |
| ECV SS | | | | 0.656 | 0.497 | 0.368 | 0.255 |
| ECV SG | | | | 0.656 | 0.106 | 0.122 | 0.116 |
| ECV GS | | | | 0.656 | 0.503 | 0.632 | 0.745 |
| Omega | | | | 0.884 | 0.755 | 0.748 | 0.826 |
| OmegaH | | | | 0.766 | 0.330 | 0.256 | 0.160 |
| H | | | | 0.938 | 0.754 | 0.679 | 0.672 |
| FD | | | | 0.957 | 0.978 | 0.844 | 0.863 |

**Table S4. Structural MRI Acquisition |** Data acquisition parameters for structural MRI data are shown for studies and acquisition sites.

| Study | Sequence | Scanner Type | Voxel Size (mm) | Image Orientation | Parallel Reduction Factor in Plane | TR (ms) | TE (ms) | Matrix Size (voxels) | Flip Angle (deg) | Dominant Group (%) |
|---|---|---|---|---|---|---|---|---|---|---|
| **Healthy Brain Network (HBN) - Staten Island** | T1 | Siemens Avanto 1.5T | 1 x 1 x 1 | LAS+ | 2 | 2730 | 1.64 | 176 x 256 x 256 | 7 | 100 |
| **Healthy Brain Network (HBN) - Rutgers University** | T1 | Siemens Tim Trio 3T | 0.8 x 0.8 x 0.8 | LAS+ | 2 | 2500 | 3.15 | 224 x 320 x 320 | 8 | 100 |
| **Healthy Brain Network (HBN) - Rutgers University** | T2 | Siemens Tim Trio 3T | 0.8 x 0.8 x 0.8 | LAS+ | 2 | 3200 | 564 | 224 x 320 x 320 | 120 | 47 |
| **Healthy Brain Network (HBN) - The City University of New York** | T1 | Siemens Tim Trio 3T | 1 x 1 x 1 | LAS+ | 2 | 2500 | 2.9 | 176 x 256 x 256 | 8 | 94 |
| **Healthy Brain Network (HBN) - The City University of New York** | T2 | Siemens Tim Trio 3T | 1 x 1 x 1 | LAS+ | 2 | 3200 | 565 | 176 x 256 x 256 | 120 | 97 |
| **Healthy Brain Network (HBN) - Citigroup Biomedical Imaging Center** | T1 | Siemens Tim Trio 3T | 1 x 1 x 1 | LAS+ | 2 | 2500 | 2.88 | 176 x 256 x 256 | 8 | 94 |
| **Healthy Brain Network (HBN) - Citigroup Biomedical Imaging Center** | T2 | Siemens Tim Trio 3T | 1 x 1 x 1 | LAS+ | 2 | 3200 | 565 | 176 x 256 x 256 | 120 | 95 |
| **Brazilian High Risk Cohort (BHRC)** | T1 | General Electric 1.5T | 0.94 x 0.94 x 1.2 | RAS+ | | 10.81 | 4.2 | 256 x 256 x 156 | 15 | 30 |
| **Philadelphia Neurodevelopmental Cohort (PNC)** | T1 | Siemens Tim Trio 3T | 0.94 x 0.94 x 1 | RAS+ | 2 | 1810 | 3.51 | 92 x 256 x 160 | 9 | 100 |
| **Nathan Kline Institute - Rockland Sample (NKI)** | T1 | Siemens Tim Trio 3T | 1 x 0.98 x 0.98 | LAS+ and/or RAS+ | 2 | 1900 | 2.52 | 176 x 256 x 256 | 9 | 45 |
| **Nathan Kline Institute - Rockland Sample (NKI)** | T2 | Siemens Tim Trio 3T | 1 x 1 x 1 | LAS+ | 2 | 3200 | 306 | 176 x 256 x 256 | 120 | 33 |
| **Developmental Component of the Chinese Color Nest Project (CCNP)** | T1 | Siemens Tim Trio 3T | 1 x 1 x 1 | RAS+ | | 2600 | 3.02 | 176 x 256 x 256 | 8 | 97 |

**Table S5. Functional MRI Acquisition |** Data acquisition parameters for structural MRI data are shown for studies and acquisition sites.

| Study | Sequence | Tasks | Total Time (min:sec) | No. of Volumes | Scanner Type | Voxel Size (mm) | TR (ms) | TE (ms) | Matrix Size (voxels) | Flip Angle (deg) | Dominant Group (%) | Field Map |
|---|---|---|---|---|---|---|---|---|---|---|---|---|
| **Healthy Brain Network (HBN) - Staten Island** | Singleband | Rest | 10:09 | 420 | Siemens Avanto 1.5T | 2.46 x 2.46 x 2.50 | 1450 | 40 | 78 x 78 x 54 | 55 | 92 | EPI |
| **Healthy Brain Network (HBN) - Rutgers University** | Multiband | Rest Peer MovieDM MovieTP | 28:43 | 2155 | Siemens Tim Trio 3T | 2.43 x 2.43 x 2.40 | 800 | 30 | 84 x 84 x 60 | 31 | 97 | EPI |
| **Healthy Brain Network (HBN) - The City University of New York** | Multiband | Rest Peer MovieDM MovieTP | 26:56 | 2020 | Siemens Tim Trio 3T | 2.43 x 2.43 x 2.40 | 800 | 30 | 84 x 84 x 60 | 31 | 97 | EPI |
| **Healthy Brain Network (HBN) - Citigroup Biomedical Imaging Center** | Multiband | Rest Peer MovieDM MovieTP | 28:43 | 2155 | Siemens Tim Trio 3T | 2.43 x 2.43 x 2.40 | 800 | 30 | 84 x 84 x 60 | 31 | 86 | EPI |
| **Brazilian High Risk Cohort (BHRC)** | Singleband | Rest | 18:00 | 540 | General Electric 1.5T | 1.88 x 1.88 x 4.5 | 2000 | 30 | 128 x 128 x 26 | 80 | 73 | None |
| **Philadelphia Neurodevelop mental Cohort (PNC)** | Singleband | Rest Frac2back ID-emo | 4:15 | 565 | Siemens Tim Trio 3T | 3 x 3 x 3 | 3000 | 32 | 64 x 64 x 46 | 90 | 93.33 | GRE |
| **Nathan Kline Institute - Rockland Sample (NKI)** | Singleband | Rest | 5:00 | 120 | Siemens Tim Trio 3T | 3 x 3 x 3.33 | 2500 | 30 | 72 x 72 x 38 | 80 | 80 | None |
| **Nathan Kline Institute - Rockland Sample (NKI)** | Multiband | Rest Checkerboard | 12:15 | 1140 | Siemens Tim Trio 3T | 3 x 3 x 3 | 645 | 30 | 74 x 74 x 40 | 60 | 29 | None |
| **Nathan Kline Institute - Rockland Sample (NKI)** | Multiband | Rest Breathhold Checkerboard | 16:03 | 688 | Siemens Tim Trio 3T | 2 x 2 x 2 | 1400 | 30 | 112 x 112 x 64 | 65 | 20 | None |
| **Developmental Component of the Chinese Color Nest Project (CCNP)** | Singleband | Rest | 15:20 | 368 | Siemens Tim Trio 3T | 3 x 3 x 3.33 | 2500 | 30 | 72 x 72 x 38 | 80 | 85 | None |

**Table S6. Quality control per imaging modality and study**

| Dataset | Structural MRI | | Functional MRI | |
|---|---|---|---|---|
| Brazil High Risk Cohort (BHRC) | Pass<br>Artifact<br>Fail | 90.52%<br>9.36%<br>0.12% | Pass<br>Fail | 97.82%<br>2.18% |
| Developmental Chinese Color Nest Project (CCNP) | Pass<br>Artifact<br>Fail | 91.28%<br>8.20%<br>0.51% | Pass<br>Fail | 98.20%<br>1.80% |
| Healthy Brain Network (HBN) | Pass<br>Artifact<br>Fail | 59.50%<br>34.93%<br>5.56% | Pass<br>Fail | 75.67%<br>24.33% |
| Nathan Kline Institute – Rockland Sample (NKI) | Pass<br>Artifact<br>Fail | 81.19%<br>16.58%<br>2.22% | Pass<br>Fail | 84.39%<br>15.60% |
| Philadelphia Neurodevelopmental Cohort (PNC) | Pass<br>Artifact<br>Fail | 90.39%<br>9.11%<br>0.50% | Pass<br>Fail | 95.24%<br>4.76% |

**Table S7. Sample size for data analysis following QC |** Structural data with "Pass" or "Artifact" labels and functional data with "Pass" labels were included in the analyses. Psychopathology analysis included individuals that had a general psychopathology score from bifactor models (i.e., p-factor).

| **Data Modality** | **Age Analysis** | | **Psychopathology Analysis** | |
|---|---|---|---|---|
| **Structural MRI** | *Before QC* | *N* = 4,827<br>(*N* = 2,061 Female; 42.7%) | *Before QC* | *N* = 4,524<br>(*N* = 1,961 Female; 43.3%) |
| | *After QC* | *N* = 4,700<br>(*N* = 2,015 Female; 42.9%) | *After QC* | *N* = 4,404<br>(*N* = 1,917 Female; 43.5%) |
| **Functional MRI** | *Before QC* | *N* = 4,614<br>(*N* = 1,976 Female; 42.8%) | *Before QC* | *N* = 4,311<br>(*N* = 1,872 Female; 43.4%) |
| | *After QC* | *N* = 4,123<br>(*N* = 1,814 Female; 44.0%) | *After QC* | *N* = 3,847<br>(*N* = 1,716 Female; 44.6%) |
